# Ethanolamine regulates CqsR quorum-sensing signaling in *Vibrio cholerae*

**DOI:** 10.1101/589390

**Authors:** Samit Watve, Kelsey Barrasso, Sarah A. Jung, Kristen J. Davis, Lisa A. Hawver, Atul Khataokar, Ryan G. Palaganas, Matthew B. Neiditch, Lark J. Perez, Wai-Leung Ng

## Abstract

The pathogen that causes cholera, *Vibrio cholerae*, uses the cell-cell communication process known as quorum sensing (QS) to regulate virulence factor production and biofilm formation in response to changes in population density and complexity. QS is mediated through the detection of extracellular chemical signals called autoinducers. Four histidine kinases, LuxPQ, CqsS, CqsR and VpsS, have been identified as receptors to activate the key QS regulator LuxO at low cell density. At high cell density, detection of autoinducers by these receptors leads to deactivation of LuxO, resulting in population-wide gene expression changes. While the cognate autoinducers that regulate the activity of CqsS and LuxQ are known, the signals that regulate CqsR have not been determined. Here we show that the common metabolite ethanolamine specifically interacts with the ligand-binding CACHE domain of CqsR *in vitro* and induces the high cell-density QS response through CqsR kinase inhibition in *V. cholerae* cells. We also identified residues in the CqsR CACHE domain important for ethanolamine detection and signal transduction. Moreover, mutations disrupting endogenous ethanolamine production in *V. cholerae* delay the onset of, but do not abolish, the high cell-density QS gene expression. Finally, we demonstrate that modulation of CqsR QS response by ethanolamine occurs inside animal hosts. Our findings suggest that *V. cholerae* uses CqsR as a dual-function receptor to integrate information from the self-made signals as well as exogenous ethanolamine as an environmental cue to modulate QS response.

**IMPORTANCE:** Many bacteria use quorum sensing to regulate cellular processes that are important for their survival and adaptation to different environments. Quorum sensing usually depends on the detection on chemical signals called autoinducers made endogenously by the bacteria. We show here ethanolamine, a common metabolite made by various bacteria and eukaryotes, can modulate the activity of one of the quorum-sensing receptors in *Vibrio cholerae*, the etiological agent of the disease cholera. Our results raise the possibility that *V. cholerae* or other quorum-sensing bacteria can combine environmental sensing and quorum sensing to control group behaviors.

## INTRODUCTION

Quorum sensing (QS) is used by a wide variety of bacteria to coordinate population-wide changes in behaviors in response to cell density (1). *Vibrio cholerae*, which causes the diarrheal disease cholera in the human host, uses QS to regulate virulence factor production, biofilm formation, Type VI secretion, metabolic regulation, and natural competence to maintain competitive fitness in various environments (2–10). Four parallel QS signaling systems pathways have been identified in *V*. *cholerae* that rely on a phosphorelay to regulate downstream gene expression (11) (Figure 1). At low cell-density (LCD), four histidine kinases CqsS, LuxPQ, CqsR, and VpsS function in parallel to phosphorylate LuxO through an intermediate phosphotransfer protein LuxU (10, 11). Phosphorylated LuxO promotes the transcription of small RNAs Qrr1-4 which promote the translation of the transcriptional regulator AphA (12, 13). Conversely, Qrr1-4 repress the translation of transcriptional regulator HapR (14, 15). Virulence and biofilm genes are expressed at LCD (4–6, 9, 11). At high cell-density (HCD), each receptor kinase detects its unique chemical messenger called autoinducer (AI) that inhibits the kinase activity upon signal binding (1). Autoinducer synthase CqsA catalyzes the production of CAI-1 (*S*-3-hydroxytridecan-4-one) which is detected by receptor CqsS (16–20) and autoinducer synthase LuxS produces AI-2 (*S*-TMHF-borate) which is detected by LuxP/Q (21–25). Thus, at HCD, when these receptors are bound to their cognate signals, kinase activity is inhibited, leading to dephosphorylation of LuxO. This prevents the transcription of Qrr 1-4 thereby inhibiting AphA production and promoting HapR translation. Type VI secretion and competence genes are expressed at HCD (3, 7). Another QS circuit was recently identified in *V. cholerae*, consisting of a cytoplasmic receptor VqmA which recognizes DPO (3,5-dimethylpyrazin-2-ol) but this system does not participate in regulating LuxO (26).

**Figure 1.**
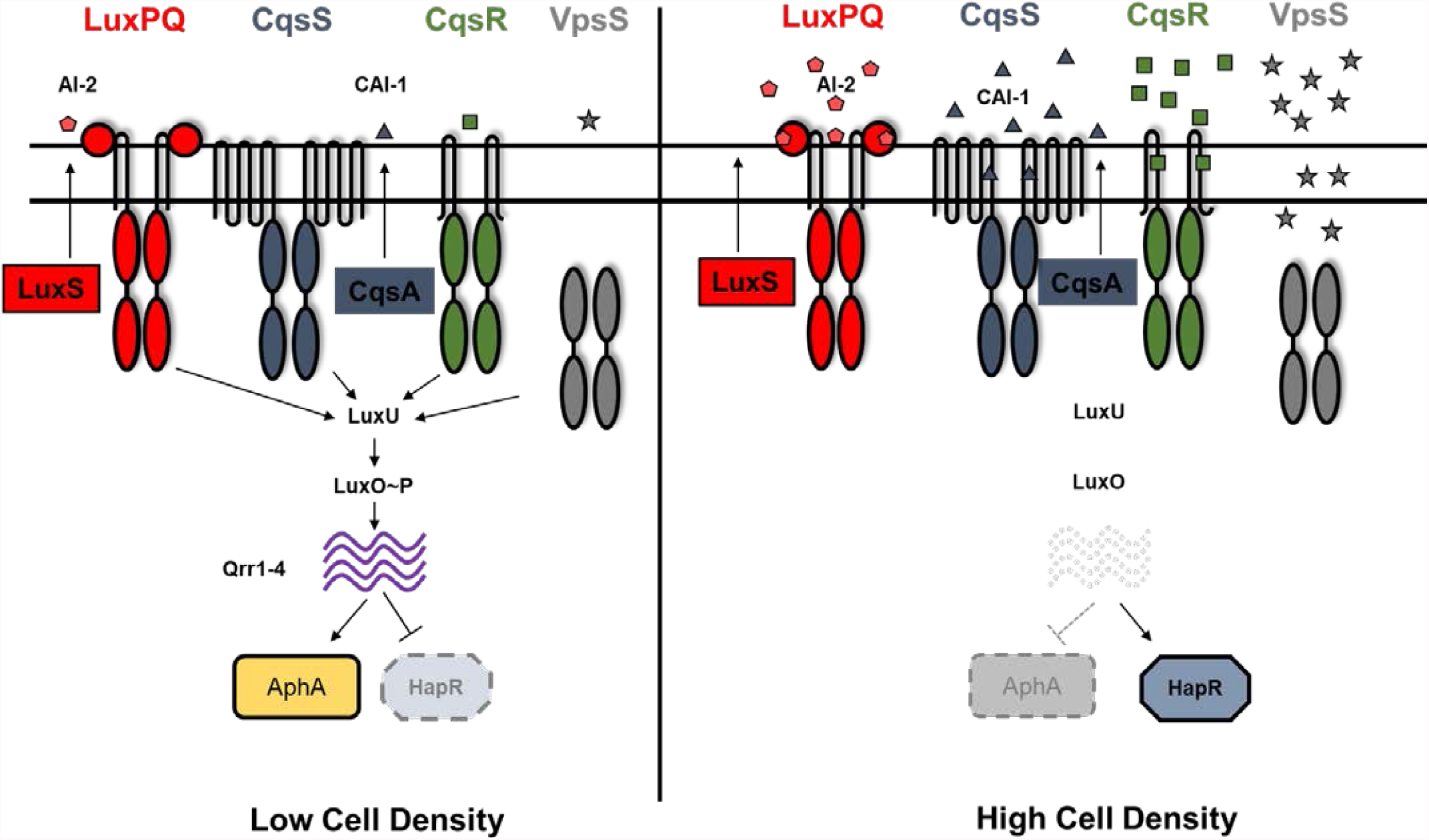
Quorum-sensing circuit in *Vibrio cholerae*. Quorum sensing in *V. cholerae* is controlled by four histidine kinases CqsS, LuxPQ, CqsR and VpsS. At low cell density, these receptors act predominantly as kinases and phosphorylate LuxO through LuxU. Phosphorylated LuxO activates transcription of small RNAs Qrr1-4 which inhibit HapR translation and promote AphA translation, thereby resulting in a low cell density expression profile. At high cell density, when the cognate signals are bound, the kinase activity of these receptors is repressed. This leads to dephosphorylation of LuxO, preventing Qrr1-4 transcription. Therefore, HapR translation is induced and AphA translation repressed, leading to a high cell density gene expression pattern. Autoinducers CAI-1 (produced by CqsA) and AI-2 (produced by LuxS) have been previously characterized in regulating the kinase activity of CqsS and LuxPQ respectively.

Although the kinase activity of both CqsR and VpsS is cell-density dependent, the signals that control the activity of CqsR and VpsS are unclear (11). VpsS has no predicted transmembrane domains and is thought to be cytoplasmic. Nitric oxide (NO) has been shown to regulate the kinase activity of VpsS *in vitro* through a signaling partner VpsV (27). However, *V. cholerae* does not make NO and Δ*vpsV* mutants has no detectable change in QS response (10, 11), so the exact role of NO sensing by VpsV in *V. cholerae* QS response remains unknown.

In contrast, CqsR is predicted to have a periplasmic CACHE domain that is known to be involved in signal sensing in many chemotaxis receptors (28). Here, by isolating constitutively active CqsR mutants, we identified several key residues within the CqsR CACHE domain required for signal binding or signal transduction. Through a chemical library screen and structure-activity relationship *in vitro* binding assay, we discovered that ethanolamine and its analogs, alaninol and serinol, bind to the CqsR CACHE domain and modulate QS response of *V. cholerae* through CqsR. We also determined that *V. cholerae* strains defective in ethanolamine biosynthesis delayed the onset of, but did not abolish, the HCD QS response. Using an infant mouse colonization model, we show that ethanolamine modulates *V. cholerae* QS response through CqsR inside animal hosts. We therefore predict that ethanolamine is used as an external cue for niche sensing and additional signals are produced by *V. cholerae* to modulate QS through CqsR.

## MATERIALS AND METHODS

### Bioinformatics

Conserved domain structures of *V. cholerae* CqsR (accession no.: NP_231465.1) were analyzed using a suite of online tools including CDD (29), CDART (30), InterPro (31), and TMHMM (32). 3D structural modelling of the CqsR ligand-binding domain was performed using HHPred (33) and Phyre2 (34) and compared to the known structure of the periplasmic region of Mlp37 (PDB ID: 3C8C) (35).

### Strains, media and culture conditions

All *V*. *cholerae* strains used in this study were derived from C6706*str*2, a streptomycin-resistant isolate of C6706 (O1 El Tor) (36). The HapR-dependent bioluminescent reporter pBB1 has been previously described (11). *V*. *cholerae* and *E*. *coli* cultures were grown with aeration in Luria-Bertani (LB) broth at 30°C and 37°C, respectively. Unless specified, media was supplemented with streptomycin (Sm, 100 μg/ml), tetracycline (Tet, 5 μg/ml), ampicillin (Amp, 100 μg/ml), kanamycin (Kan, 100 μg/ml), chloramphenicol (Cm, 5 μg/ml) and polymyxin B (Pb, 50 U/ml) when appropriate. Bacterial strains used in this study is provided in Supplementary Table 1.

### DNA manipulations and strain construction

All DNA manipulations were performed using standard procedures. Deletions and point mutations were introduced into the *V*. *cholerae* genome by allelic exchange using the suicide vector pKAS32 or a modified version, pJT961, as previously described (11, 37, 38). Mutant strains carrying the desired mutations were screened and confirmed by PCR and sequencing.

### Bioluminescence assays

Bioluminescence assays were performed as described previously (11) with slight modifications. For manual assays, single colonies were grown overnight at 30 °C in LB containing appropriate antibiotics and the cultures were diluted at least 100-fold in the same medium. Diluted cultures were grown at 30 °C and OD_600_ (1 ml of culture) and light production (0.2 ml of culture) were measured every 45–60 min until OD_600_ reached ∼2.0 using a Thermo Scientific Evolution 201 UV-Visible Spectrophotometer and a BioTek Synergy HT Plate Reader, respectively. For automated assays, single colonies were grown in LB medium containing appropriate antibiotics in 96- or 384-well microplates at 30°C with aeration. OD_600_ and light production were measured every 30 min for at least 10 hr using a BioTek Synergy HT Plate Reader. Light production per cell was calculated from dividing light production by OD_600_.

### Random mutagenesis and screening of constitutively active CqsR mutants

The region encoding the predicted ligand-binding domain of CqsR (amino acids 31-297) was amplified from pEVS143-*cqsR* (11, 39) with primers WNTP0548 (GAGTTATTGGGGGCTTGAAGT) and WNTP0549 (AATCCTTTTTCGATGTTGATAATTAAGTCG) and subjected to random mutagenesis using GeneMorph II EZClone Mutagenesis kit (Agilent Technologies) per manufacturer’s instructions. The resulting *cqsR* random mutagenized library was transformed into XL10-Gold *E. coli* cells. A portion of the library was then conjugated into a quadruple QS receptor strain (Δ4, Δ*cqsS*, Δ*luxQ*, Δ*cqsR*, Δ*vpsS*) carrying a HapR-dependent bioluminescence reporter pBB1 (11) and the transconjugants were selected on LB/Pb/Kan/Tet2 plates and used for screening of constitutively active CqsR mutants. Colonies were picked and transferred into microplates containing LB/Kan/Tet2/IPTG medium using a colony picking robot and grown at 30°C overnight. OD_600_ and luminescence were measured with a BioTek Synergy HT Plate Reader. Mutants of interest that displayed low relative bioluminescence were collected, re-screened again to confirm. The mutations in the plasmid-borne *cqsR* of these isolates were determined by sequencing. To introduce specific amino acid changes, site directed mutagenesis on the plasmid-borne *cqsR* was performed using the QuikChange™ XL Site-Directed Mutagenesis Kit (Agilent) per manufacturer’s instructions.

### NMR metabolomics

Extracellular metabolite analysis of *V. cholerae* culture supernatants was performed as described previously (8) except plain LB medium was used as reference standard. An ^1^H NMR spectrum of each sample was collected at 25°C on a Bruker Avance 600 spectrometer by using 64 scans and a NOE1D pulse sequence. Data were processed and analyzed by using CHENOMX (version 8.0) for quantification of metabolites present in each sample. The average value and the standard error of the mean (SEM) were determined for at least three replicates.

### CqsR LBD purification

The region encoding the periplasmic region of CqsR (CqsR-LBD, residues 35-274) was PCR amplified with primers WNTP0694 (GCCTGGTGCCGCGCGGCAGCGAAGTCCCATTTAGAAAAGAG) and WNTP695 (GCTTTGTTAGCAGCCGGATCTTAGATGTTGATAATTAAGTCGAAG) from *V. cholerae* genomic DNA; the vector pET28B was amplified with primers WNTP692 (GCTGCCGCGCGGCACCAG) and WNTP693 (GATCCGGCTGCTAACAAAG). The two fragments were joined together using Gibson reaction (NEB) and the resulting plasmid was transformed into *E. coli* BL21(DE3) strain (WN5327).

For CqsR-LBD purification, strain WN5327 was grown to OD_600_ ∼ 0.5 and induced with 1 mM IPTG at 16 °C overnight. Cells were collected by centrifugation and resuspended in binding buffer (50mM sodium phosphate buffer, pH 7.0; 300mM sodium chloride; 10mM imidazole, pH 7.7; 5% glycerol). Resuspended cells were then lysed with a fluidizer. Insoluble materials were removed by centrifugation (10,000g, 4 °C, 1 hour) and subsequent filtering through a 0.45 µm filter. Cleared lysate was loaded onto His-Trap column (1 mL) equilibrated with binding buffer. The column was then washed with 20 mL binding buffer. Proteins were eluted with binding buffer containing 120 to 300 mM imidazole. Fractions containing CqsR-LBD were pooled, frozen in aliquots, and stored at −80°C.

### Differential scanning fluorimetry

Differential scanning fluorimetry assays were performed using a BioRad CFX Connect Real-Time PCR instrument (40). For chemical screening, ligands were prepared by dissolving the contents of Biolog plates PM 1-4 in 50 μl of water (final concentration ∼10–20 mM). Each 20 μl standard assay contained 1× PBS (pH 7.5), 10%(v/v) glycerol, 10 μM CqsR-LBD, and 5× SYPRO Orange. Two μl of the resuspended Biolog compounds were added to each assay. Samples were heat denatured from 20°C to 90°C at a ramp rate of 1°C min^−1^. The protein unfolding curves were monitored by detecting changes in SYPRO Orange fluorescence. The first derivative values (−dF/dt) from the raw fluorescence data were used to determine the melting temperature (Tm). For testing other compounds, a 10× stock was prepared and 2 μl was added to each well and the experiments were conducted as described above.

### MicroScale Thermophoresis (MST)

Nickel affinity resin-purified CqsR-LBD was further purified using anion exchange chromatography (Source15Q - GE Healthcare) and size exclusion chromatography (S200 – GE Healthcare). Purified CqsR-LBD was concentrated to 4.3 mg/ml and stored in 50 mM sodium phosphate, pH 7.7, and 300 mM sodium chloride. CqsR-LBD was diluted to 20 μM and labeled using the Monolith NT Protein Labeling Ki RED-NHS (NanoTemper Technologies). Different concentrations of ethanolamine were incubated with 20 nM working stock solutions of labeled protein in the dark for 30 min at 4° C. After incubation, the samples were transferred into standard treated capillaries (NanoTemper Technologies) and read in a Monolith NT.115 Blue/Red instrument at room temperature using 20% LED and medium MST power. Binding affinities were calculated from three experiments.

### Chemical Synthesis of EA Metabolites

See supplementary information for chemical synthesis of HEGly and HEHEAA.

### Ethics statement

All animal experiments were done in accordance with NIH guidelines, the Animal Welfare Act, and US federal law. The infant mouse colonization experimental protocol B-2018-99 was approved by Tufts University School of Medicine’s Institutional Animal Care and Use Committee. All animals were housed in a centralized and AAALAC-accredited research animal facility that is fully staffed with trained husbandry, technical, and veterinary personnel in accordance with the regulations of the Comparative Medicine Services at Tufts University School of Medicine.

### Infant mouse colonization model

For *V. cholerae* infection, bacterial strains were grown aerobically overnight in LB/Sm at 30°C and diluted in LB for inoculum. Approximately 10^6^ colony forming units (CFUs) in 50 µl were introduced orogastrically to 3- to 5-day-old CD-1 mice (Charles River Laboratories). Infected animals were subsequently gavaged with 50µL 10mM ethanolamine (EA) or vehicle (LB) at 2 and 4 hr post-infection. All animals were sacrificed 8 hours after infection and their small intestines were harvested and homogenized. *V. cholerae* colonized in the small intestine was enumerated by plating serial dilutions of intestinal homogenate on LB/Sm plates. *V*. *cholerae* colonization of the small intestine is presented as a single data point per mouse and data are graphed with the median. Two-tailed unpaired Student’s t tests, assuming unequal variances were used for statistical analyses **P < 0.01.

## RESULTS

### CqsR is predicted to carry a periplasmic CACHE domain

Similar to other hybrid histidine kinases, CqsR possesses a cytoplasmic phosphoacceptor domain (aa 355 – 421), an HATPase domain (aa 363 – 583) and a C-terminal response regulator domain (aa 603-720). Using various bioinformatics tools, the periplasmic region of CqsR is predicted to carry a periplasmic dCACHE_1 domain ((28, 41), aa 44-239, simplified as CACHE domain hereafter) (Figure 2A). The predicted structure contains the characteristic long N-terminal α-helix (green), a linker (yellow) that connects this helix to two globular domains (pocket I, red and pocket II, blue) with a small α-helical linker between the two pockets (orange), and a C-terminal α-helix (purple) that connects the domain to the cytoplasm (Figure 2B), strikingly similar to other CACHE proteins, such as *V. cholerae* chemoreceptor Mlp37 (35) (Figure 2B). These similarities strongly suggest that the periplasmic region of CqsR functions as a ligand-binding domain to regulate its cytoplasmic kinase activity.

**Figure 2.**
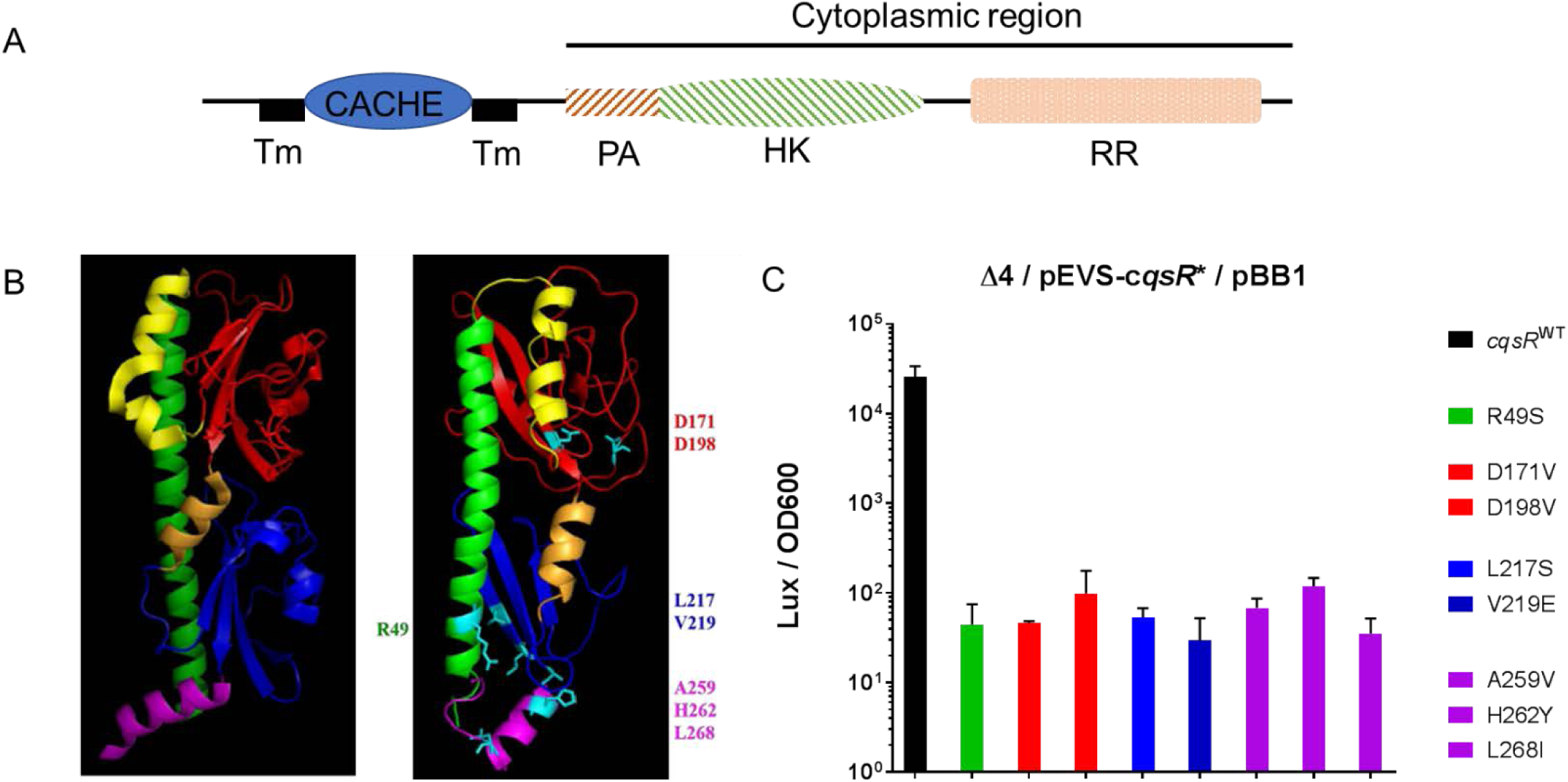
The CqsR CACHE domain is important for signal sensing and signal transduction. (A) CqsR is predicted to contain all the conserved cytoplasmic domains (phosphoacceptor (PA), histidine kinase (HK), and response regulator (RR) domains) of a hybrid histidine kinase with an N-terminal periplasmic CACHE domain flanked by two transmembrane helices (TM). (B) The predicted structure of the periplasmic CACHE domain of CqsR (right) is similar to that of Mlp37 (left). The residues identified as important for signal sensing and signal transduction are highlighted in cyan in the predicted structure. See main text for further details. (C) Eight periplasmic CqsR residues are involved in signal sensing and signal transduction. Relative light production (lux/OD_600_) was measured in quadruple QS receptor (Δ4) strains carrying a plasmid producing CqsR with single amino changes as well as a HapR-dependent bioluminescence reporter. Alteration in the CqsR helix domain (R49S), CACHE pocket 1 region (D171V, D198V), CACHE pocket 2 region (L217S, V219E), or the transmembrane proximal regions (A259V, H262Y, L268I) impaired bioluminescence production at high cell-density (OD_600_ > 1.5). Average values and standard errors from at least three independent replicates are shown.

### Identification of residues in the CqsR periplasmic region important for signal sensing or signal transduction

CqsR kinase activity is inhibited by an uncharacterized activity present in the spent culture media from *V. cholerae* (11). We therefore hypothesized that CqsR detects a discrete chemical signal through the periplasmic CACHE domain and predicted that certain mutations in this domain could impair signal sensing and signal transduction. Consequently, these mutations would turn CqsR into a constitutively active kinase even in the presence of the cognate signals. To identify these mutations, we constructed a *cqsR* plasmid library in which the whole periplasmic region was randomly mutagenized. The randomly mutagenized *cqsR* library was then introduced into a quadruple receptor mutant (Δ4) *V. cholerae* strain (Δ*luxQ*, Δ*cqsS*, Δ*vpsS*, Δ*cqsR*) that also harbors a HapR-dependent bioluminescent reporter cosmid pBB1 (11). While the quadruple receptor mutant produces light constitutively due to the constant production of HapR, in the presence of a plasmid-borne wild type (WT) copy of *cqsR*, this strain produces low bioluminescence at low cell-density (LCD) and subsequently turns on bioluminescence production at high cell-density (HCD) due to signal detection and kinase inhibition of CqsR (11). Thus, we predicted that clones carrying plasmids expressing WT *cqsR* or a null *cqsR* allele would appear bright at HCD, while clones producing CqsR variants insensitive to the presence of cognate signal would appear darker at HCD. We measured the bioluminescence of ∼26,000 clones from this random library and identified 79 candidates impaired for light production at HCD. Most candidates harbored multiple mutations in the plasmid-borne *cqsR*. There were 23 unique mutations in these candidates. When each unique mutation was introduced back to *cqsR* individually, only eight unique mutations in *cqsR* resulted in lower bioluminescence production at HCD (Figure 2C). These *cqsR* mutations caused changes in the two aspartate residues (D171V and D178V) mapped to pocket I of the predicted ligand binding domain, two changes in pocket II (L217S and V219E), four changes (R49S, A259V, H262Y, and L268I) in the two helical regions that either exit or enter the cellular membrane and are likely not involved in direct ligand interactions. Strikingly, both Asp171 and Asp198 residues in the pocket I of CqsR are predicted to be in the same positions as the Asp172 and Asp201 residues respectively, in Mlp37, which have been shown to be critical for binding to amino acid signals (35). Therefore, our results suggest that, similar to Mlp37, CqsR detects its cognate signals through the periplasmic CACHE domain.

### Chemical screen identified ethanolamine as a potential CqsR ligand

Since the periplasmic CACHE domain is critical for the signal sensing, we attempted to identify potential ligands capable of binding to the putative CqsR ligand-binding domain (CqsR-LBD), by performing a differential scanning fluorimetry screen using the common metabolites contained within the Biolog PM1-4 plates. This assay assumed that ligand binding leads to stabilization of the target protein, resulting in an increase of melting temperature (Tm) of the protein, which can be determined by measuring the dynamics of Sypro Orange fluorescence emission at different temperatures. In the absence of exogenous metabolites, the average Tm of the purified CqsR-LBD was observed to be 46 °C under the test conditions. Out of the ∼400 metabolites tested, including some of the known compounds that bind CACHE domain such as amino acids, carboxylates, and polyamines, we only observed a shift in the Tm of >5 °C in the presence of one tested compound, ethanolamine (or 2-aminoethanol). The shift in Tm of CqsR in the presence of ethanolamine was dose-dependent. The largest shift in Tm of +15 °C was observed for 10 mM ethanolamine (Table 1) with smaller shifts of +9.33 °C, +6°C and +2°C for 1 mM, 100 µM and 10 µM ethanolamine respectively. To investigate the specificity of the interactions between ethanolamine and the CqsR-LBD, we determined how chemical modifications to the ligand affected CqsR-LBD binding (Table 1). Replacing the hydroxyl group of ethanolamine with a thiol, methyl, or carboxyl group abolished *in vitro* binding to CqsR-LBD, as indicated by the lack of change in the observed Tm in the presence of these compounds. Similarly, replacing the amine group with a thiol or methyl group also abolished *in vitro* binding (Table 1). In addition, single or multiple methylations of the amine group of ethanolamine also abolished binding (Table 1). We therefore concluded that the presence of both the amine and the hydroxyl groups in ethanolamine are essential for efficient CqsR-LBD binding. Side-chain substitutions at the carbon atom next to the hydroxyl group (labelled i in Table 1) also abolished CqsR-LBD binding regardless of the stereochemistry of the hydroxyl group. However, substitutions on the carbon atom next to the amine group (labelled ii) appeared to be better tolerated, as indicated by the increase in Tm observed for L-alaninol and serinol (Table 1). Interestingly, stereospecificity is critical for binding activity as indicated by lower shift in Tm for D-alaninol, a stereo-isomer of L-alaninol, at the same concentration (Table 1).

**TABLE 1.**
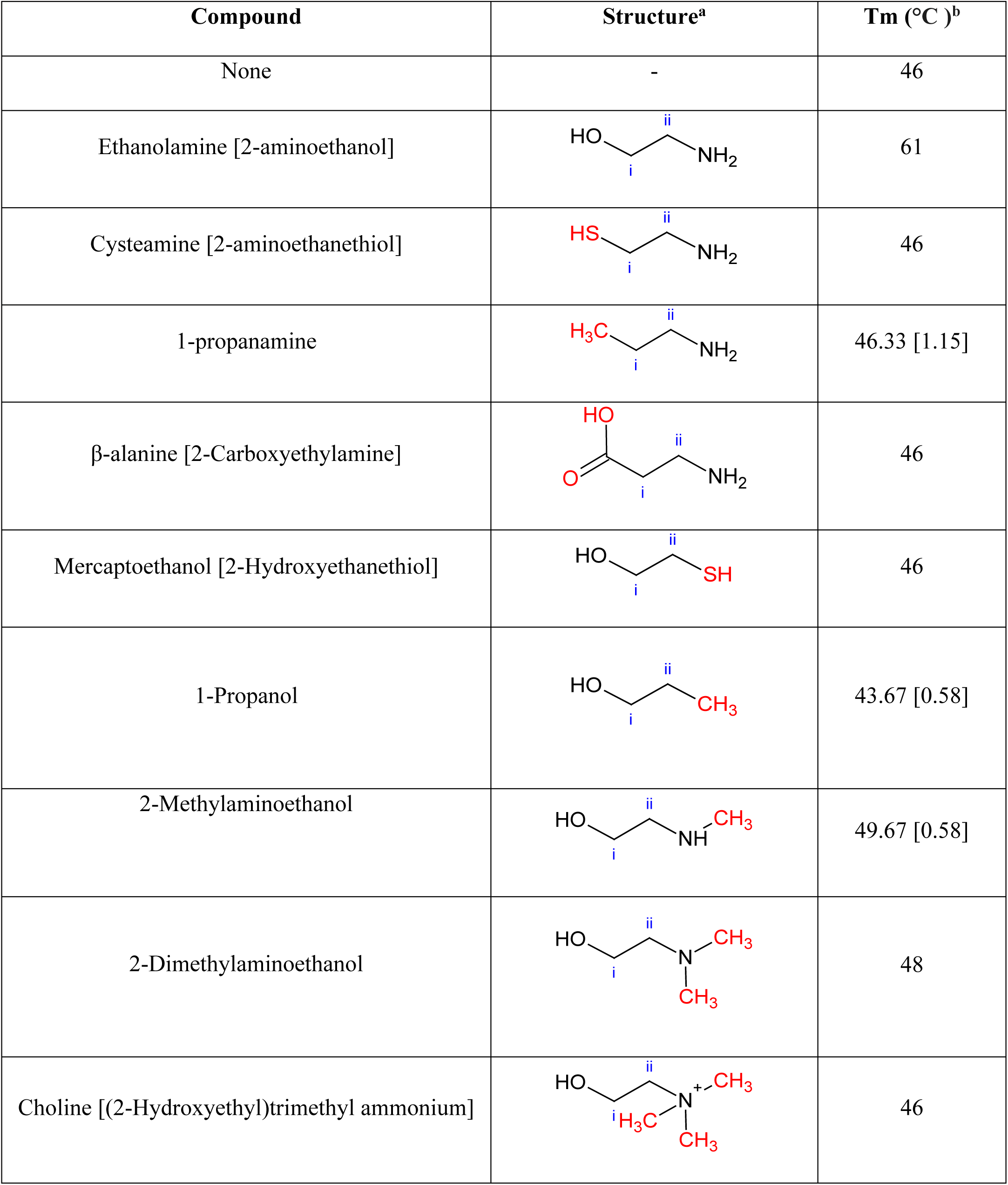
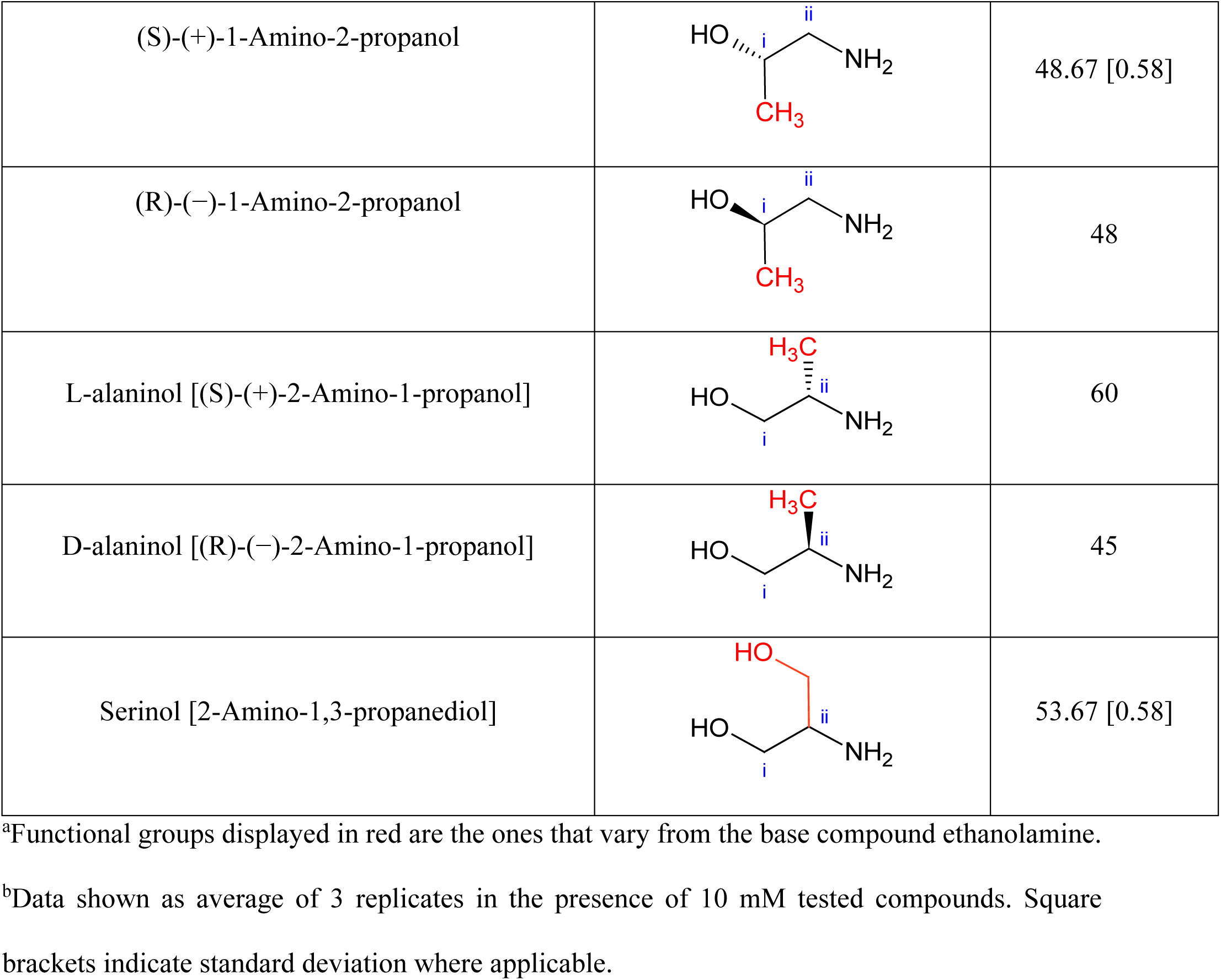
Effects of ethanolamine and its analogs on the melting temp of CqsR-LBD.

| Compound | Structure<sup>a</sup> | T<sub>m</sub> (°C)<sup>b</sup> |
| --- | --- | --- |
| None | - | 46 |
| Ethanolamine [2-aminoethanol] |  | 61 |
| Cysteamine [2-aminoethanethiol] |  | 46 |
| 1-propanamine |  | 46.33 [1.15] |
| β-alanine [2-Carboxyethylamine] |  | 46 |
| Mercaptoethanol [2-Hydroxyethanethiol] |  | 46 |
| 1-Propanol |  | 43.67 [0.58] |
| 2-Methylaminoethanol |  | 49.67 [0.58] |
| 2-Dimethylaminoethanol |  | 48 |
| Choline [(2-Hydroxyethyl)trimethyl ammonium] |  | 46 |
| (S)-(+)-1-Amino-2-propanol |  | 48.67 [0.58] |
| (R)-(-)-1-Amino-2-propanol |  | 48 |
| L-alaninol [(S)-(+)-2-Amino-1-propanol] |  | 60 |
| D-alaninol [(R)-(-)-2-Amino-1-propanol] |  | 45 |
| Serinol [2-Amino-1,3-propanediol] |  | 53.67 [0.58] |

<sup>a</sup>Functional groups displayed in red are the ones that vary from the base compound ethanolamine.
<sup>b</sup>Data shown as average of 3 replicates in the presence of 10 mM tested compounds. Square brackets indicate standard deviation where applicable.

A recent study showed that a specific ethanolamine derivative, N-(2-hydroxyethyl)-2-(2-hydroxyethylamino) acetamide [HEHEAA], present in commercial preparations of ethanolamine, is responsible for promotion of *pipA* transcription in *Pseudomonas sp. GM79* in response to plant exudates (42). We tested eight common compounds that accumulate during chemical synthesis of ethanolamine including HEHEAA (42, 43) for their ability to bind to CqsR and, in our hands, none of the compounds tested produced a change in Tm of the CqsR-LBD in the binding assay (Supplementary Table 2), suggesting the CqsR response to ethanolamine was not due to these common contaminants. Finally, using MicroScale Thermophoresis, we determined the binding affinity (Kd) of ethanolamine to CqsR-LBD is ∼0.5 µM (Supplementary Figure 1). Collectively, our data indicate that ethanolamine and some ethanolamine derivatives bind to CqsR-LBD with a high degree of specificity.

### Effect of ethanolamine and its analogs in regulating *V. cholerae* quorum sensing

Since ethanolamine binds to the CqsR-LBD *in vitro*, we postulated that exogenous addition of ethanolamine might diminish CqsR kinase activity, leading to premature induction of HCD QS response in *V. cholerae* cells. To test this, we measured the density-dependent bioluminescence profile of a Δ3 *cqsR*^+^ (Δl*uxQ*, Δ*cqsS*, Δ*vpsS*) strain that harbors the HapR-dependent bioluminescent reporter pBB1 in the presence of varying concentrations of exogenously added ethanolamine. In the absence of ethanolamine, similar to previously reported (11), the Δ3 *cqsR*^+^ strain displayed the characteristic U-shaped HapR-dependent bioluminescence profile due to changes in CqsR kinase activity in different cell densities (Figure 3A). Induction of HCD QS response by ethanolamine, as observed by increased levels of light production at low cell densities, was dose-dependent (Figure 3A). In the presence of 10 mM exogenously added ethanolamine, the strain was constitutively bright at all cell densities when compared to that from the culture in LB medium without ethanolamine added (Figure 3A). Moreover, lower concentrations (1mM or 0.1mM) of exogenous ethanolamine also induced a higher level of light production in this reporter strain. Consistent with our model, using a P*qrr*4-*lux* reporter (11), we also determined that ethanolamine repressed Qrr4 transcription in the Δ3 *cqsR*^+^ strain (Supplementary Figure 2). Similar QS-inducing effects were obtained for L-alaninol and serinol (Figure 3B, 3C and Supplementary Figure 2). Consistent with the observation that D-alaninol did not show any detectable *in vitro* binding activity to CqsR, 10 mM D-alaninol was inactive in inducing QS gene expression (Table I, Figure 3D, and Supplementary Figure 2). In addition, ethanolamine did not induce HapR-dependent bioluminescence in other triple QS receptor mutants where only CqsS, LuxPQ, or VpsS is present as the sole QS receptor (Supplementary Figure 3), indicating that ethanolamine induces QS in a CqsR specific manner. Finally, unlike the strain producing WT CqsR, 10 mM of ethanolamine did not induce constitutive light production in the CqsR^D171V^ and CqsR^D198V^ CACHE domain mutants (Figure 3E and Figure 3F). Together, our results strongly suggest that ethanolamine specifically interacts with the CACHE domain of CqsR to inhibit its kinase activity and modulate QS gene expression in *V. cholerae*.

**Figure 3.**
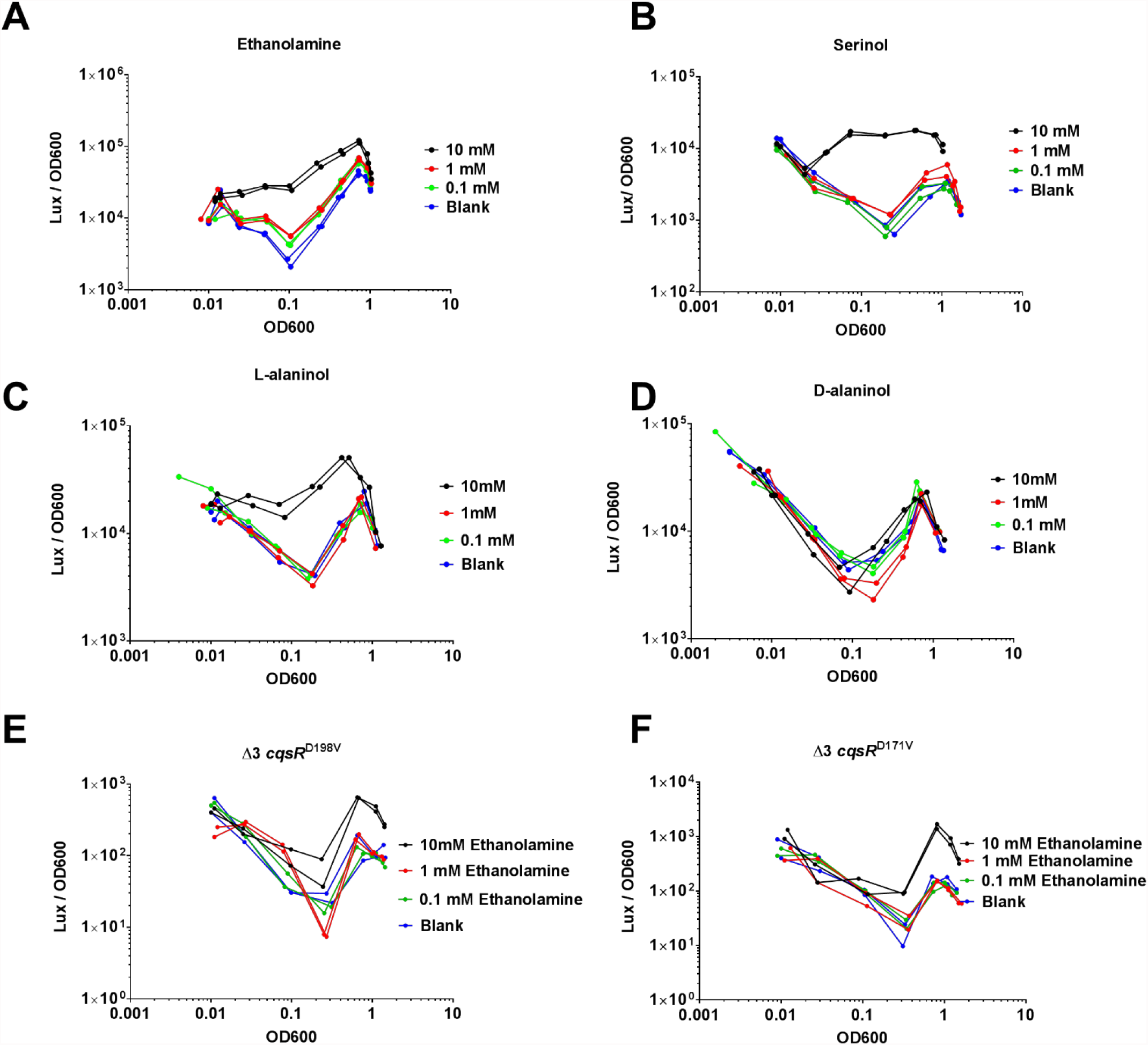
Effects of ethanolamine and its analogs on CqsR quorum-sensing response. HapR-dependent bioluminescence profiles (lux/OD_600_) were measured in a Δ3 *cqsR*^+^ strain in the presence of 10 mM, 1mM, 0.1 mM A) ethanolamine, B) serinol, C) L-alaninol, D) D-alaninol. Blank indicates LB medium without ethanolamine added. HapR-dependent bioluminescence profiles (lux/OD_600_) were measured in E) Δ3 *cqsR*^D198V^ strain and F) Δ3 *cqsR*^D171V^ strain in the presence of 10 mM, 1mM, 0.1 mM concentrations ethanolamine respectively. Each figure shows a representative profile of each condition with two biological replicates. Each experiment was performed independently at least two times.

### The role of intrinsically produced ethanolamine in QS

We showed above that exogenous ethanolamine inhibits CqsR kinase activity in *V. cholerae*, we then tested if endogenously made ethanolamine also plays a role in controlling CqsR. In *E. coli*, ethanolamine is derived from the degradation of glycerol-3-phosphoethanolamine to glycerol-3-phosphate and ethanolamine (44, 45). This reaction is predicted to be catalyzed by two *V. cholerae* glycerophosphodiester phosphodiesterases VCA0136 (GlpQ homolog) and VC1554 (UgpQ homolog). We used ^1^H NMR to determine the ethanolamine level in the cell-free supernatants derived from both the Δ3 *cqsR*^+^ and the Δ3 *cqsR*^+^ Δ*vca0136* Δ*vc1554* strains. We detected ∼25 µM ethanolamine in the supernatants harvested from the Δ3 *cqsR*^+^ strain but the level of ethanolamine was below the detection limit in the supernatants from the Δ3 *cqsR*^+^ Δ*vca0136* Δ*vc1554* strain.

After confirming the double phosphodiesterase mutants did not make any ethanolamine, we used the HapR-dependent bioluminescence pBB1 reporter to measure the QS response of these strains. We expected the mutant is impaired in expressing the HCD QS response due to the lack of ethanolamine to repress the CqsR kinase activity. However, the double phosphodiesterase mutant still exhibited density-dependent U-shaped bioluminescence profile similar to the Δ3 *cqsR*^+^ strain. Nevertheless, the onset of the transition to HCD bioluminescence production was delayed in the double phosphodiesterase mutants (Figure 4), suggesting the inhibition of CqsR kinase activity and HapR production occur at a higher cell density. Consistent with the above results, using a P*qrr*4-*lux* reporter, light production at LCD was higher in the Δ3 *cqsR*^+^ Δ*vca0136* Δ*vc1554* strain when compared to that of the Δ3 *cqsR*^+^ strain (Supplementary Figure 4), indicating transcription of Qrr sRNAs is increased in the absence of endogenous ethanolamine production in the double phosphodiesterase mutants, resulting in a delayed HapR production. It should be noted that, even in the absence of endogenous ethanolamine production, repression of Qrr transcription and induction of HapR production still occur, indicating the presence of additional signals inhibiting CqsR kinase activity at HCD (Figure 4 and Supplementary figure 4). Moreover, addition of 10 mM ethanolamine increased light production in Δ3 *cqsR*^+^ as well as in the Δ3 *cqsR*^+^ Δ*vca0136* Δ*vc1554* carrying the pBB1 HapR-dependent reporters (Figure 4) and decreased light production in the same strains carrying the P*qrr*4-*lux* reporters (Supplementary figure 4), indicating CqsR sensing is functional in responding to exogenously supplied ethanolamine in these strains. Together, our results indicate that while endogenously made ethanolamine participates in QS gene regulation, *V. cholerae* produces additional signals that are capable of activating QS through CqsR.

**Figure 4.**
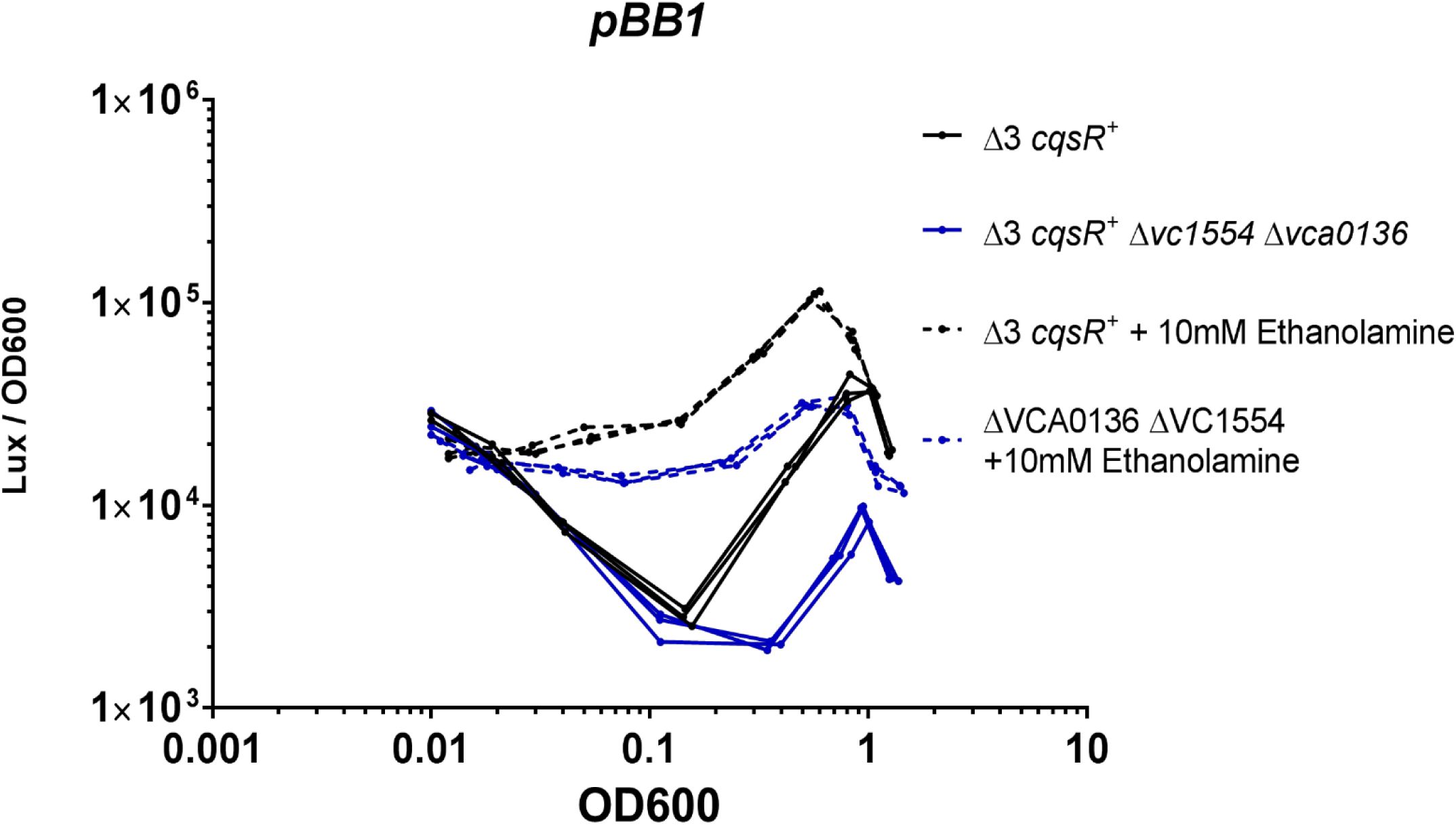
Effect of glycerophosphodiesterase deletions on CqsR quorum-sensing response. HapR-dependent bioluminescence profiles (lux/OD_600_) were measured in a Δ3 *cqsR*^+^ strain and an isogenic strain with deletions in loci *vc1554* and *vca0136*, encoding the two putative glycerophosphodiester phosphodiesterases in LB medium and in the presence of 10 mM ethanolamine. The figure shows a representative profile of each condition with two biological replicates. Each experiment was performed independently at least two times.

### Ethanolamine modulates CqsR QS signaling inside animal hosts

Production of the Toxin-Coregulated Pili, which is critical for *V. cholerae* host colonization, is repressed by the HCD QS response (5, 6). We previously showed that the Δ3 *cqsR*^+^ strain, but not the Δ4 strain, can colonize the small intestines of infant mice (11). To determine if exogenous ethanolamine influences CqsR signaling *in vivo*, we assayed the ability of the Δ3 *cqsR*^+^ strains to colonize the host in the presence of exogenous ethanolamine. While the Δ3 *cqsR*^+^ strain showed a ∼5-fold decrease in mouse small intestine colonization in the presence of ethanolamine compared to the group given the vehicle control, the Δ3 *cqsR*^D171V^ strain showed no significant impairment in the ability to colonize the infant mouse in the presence of ethanolamine (Figure 5). These results indicate that high levels of ethanolamine in the gut can prematurely induce the HCD QS response in the Δ3 *cqsR*^+^ strain and decrease colonization. Importantly, the effect of ethanolamine is dependent on the interaction between the compound and the CACHE domain of the CqsR.

**Figure 5.**
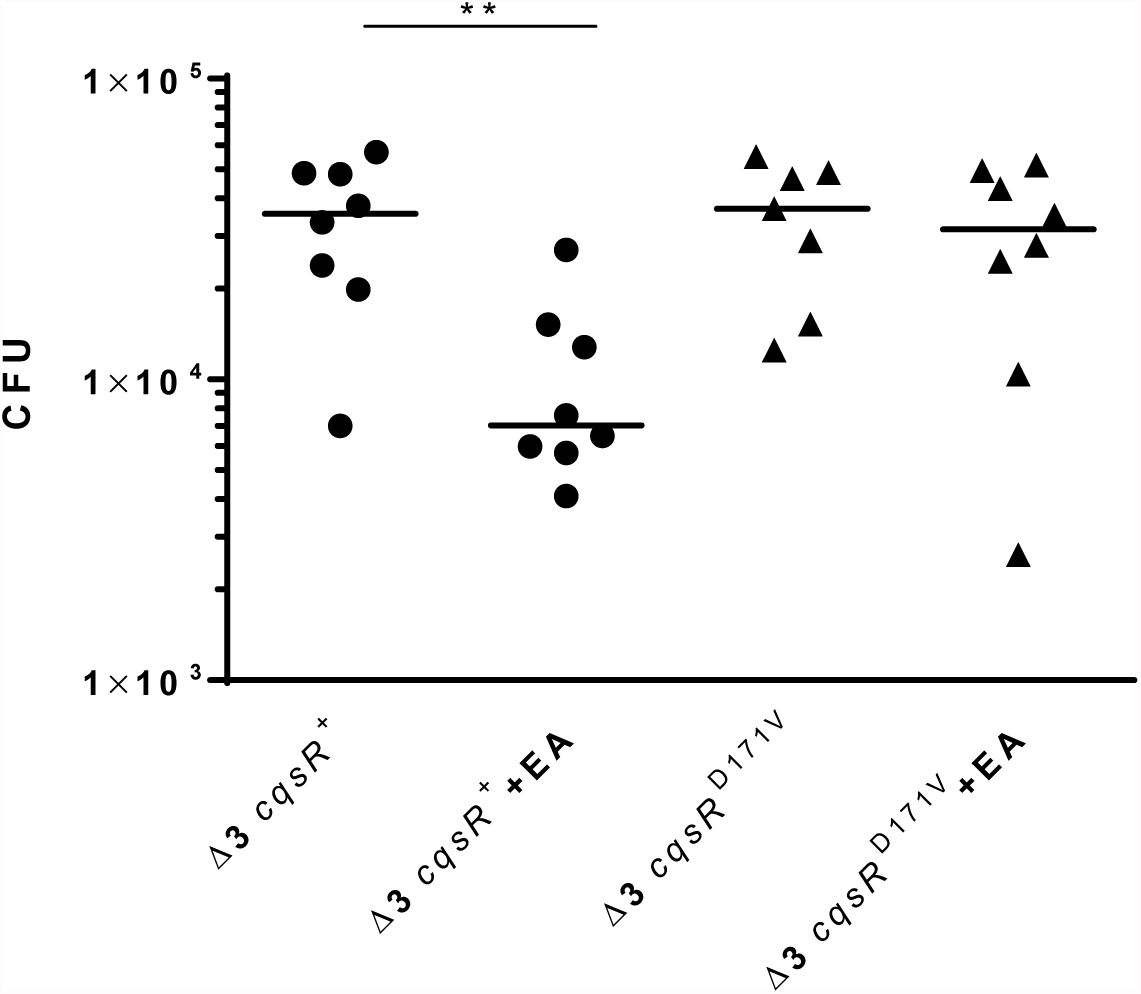
Effect of exogenous ethanolamine on CqsR quorum-sensing response inside animal hosts. CFU counts per small intestine homogenate collected from mice (n = 7-8) infected with a Δ3 *cqsR*^+^ or Δ3 *cqsR*^D171V^ mutant strain. To assess the effect of ethanolamine on strain colonization, mice were gavaged with 10mM ethanolamine (EA) or vehicle at 2 and 4 hours post infection. Each symbol represents the CFU enumerated from an individual mouse and the horizontal lines indicate the median for each group. **P < 0.01 (unpaired t test).

## DISCUSSION

In this study, we demonstrated that the periplasmic CACHE domain of CqsR QS receptor is involved in signal sensing; and we also identified a common metabolite ethanolamine as a signal that regulates the CqsR QS signaling pathway in *V. cholerae*. The periplasmic CACHE domain is common in many chemotaxis receptors and other membrane bound regulators (28); and CACHE domain proteins have been shown to interact with a variety of chemical compounds (28, 35, 41, 46–48). While CACHE domain has been shown to interact with quaternary ammonium including choline (an ethanolamine analog that CqsR does not interact) to control chemotaxis (49, 50), our study provided an example where a CACHE domain interacts with an amino alcohol (i.e., ethanolamine) to regulate QS gene expression. Of note, the CACHE domain of CqsR shares no homology to that of another ethanolamine-sensing histidine kinase HK17 (EutW) in *Enterococcus faecalis* or the cytoplasmic transcriptional regulator EutR that detects ethanolamine in *Salmonella* (51, 52).

Although ethanolamine binds to CqsR-LBD with sub-micromolar affinity (k_d_ ∼0.5 µM), millimolar level of ethanolamine is needed to fully induce the HCD QS response in *V. cholerae*. This apparent difference is not uncommon for signaling pathways dependent on CACHE receptor proteins (e.g.,(50, 53)). This is in stark contrast to the typical effective concentration for canonical autoinducers, such as CAI-1 and AI-2, which are in micromolar levels (16–19, 24, 25). Nonetheless, interaction of ethanolamine to CqsR is highly specific and modifications of almost any functional groups in ethanolamine result in total loss of activity. We reasoned that while ethanolamine binds to CqsR-LBD with high affinity *in vitro*, the binding affinity to the full length CqsR inside *V. cholerae* cells could be different. Moreover, the import mechanism of ethanolamine into the *V. cholerae* periplasmic space is unclear. Indeed, *V. cholerae* is not known to metabolize ethanolamine as either carbon or nitrogen source (54) and it lacks the known ethanolamine transporters for importing the compound into the cytoplasm (55, 56). In addition, since the mutants defective in ethanolamine synthesis are still able to express a delayed HCD QS response, additional signals must be present and detected by CqsR, and these molecules could compete with ethanolamine for CqsR binding.

Given *V. cholerae* does not use ethanolamine as a nutrient source, it is unclear why CqsR detects this specific metabolite out of many secreted molecules. However, ethanolamine has been implicated in niche recognition in several pathogens such as enterohaemorrhagic *E. coli* (EHEC), *S. typhimurium* (57), *E. faecalis* (58), and *Clostridium difficile* (59). Our results suggest that ethanolamine signaling may only operates in environments where ethanolamine is found in high abundance. Because ethanolamine is derived from the membrane phospholipid turnover, this compound is particularly prevalent in the gastrointestinal tract, where it can be found at concentrations over 2 mM per some reports (60). Thus, ethanolamine might act primarily as an environmental cue as opposed to a *bona fide* QS signal in *V. cholerae*. In *Salmonella*, ethanolamine metabolism genes and Type 3 secretion system genes are activated with high and low level of ethanolamine, respectively (57, 61). Similarly, in EHEC, exogenous ethanolamine increases the expression of virulence genes (51). However, unlike these examples in which ethanolamine sensing promotes virulence gene expression, ethanolamine detection through CqsR instead represses virulence gene expression by promoting HCD QS response in *V. cholerae* (11). This is especially relevant to the life cycle of *V. cholerae* since the major colonization site of this pathogen is the small intestine while ethanolamine may be more abundant in the lower gastrointestinal (GI) tract because it cannot be metabolized by the majority of commensal species (62), thus, ethanolamine sensing may prevent *V. cholerae* from colonizing in an undesirable niche.

It is puzzling why *V. cholerae* QS circuit is composed of four functionally-redundant pathways, we previously suggest that the “many-to-one” arrangement is important for the robustness of the system to prevent signal perturbation and the HCD QS response is only expressed when all four receptors are bound with the cognate signals (11). Our results here suggest that the four QS receptors integrate information derived from different sources. It has been suggested CAI-1 is used for inter-*Vibrio* signaling (16, 18) and AI-2 is for inter-species signaling (63, 64); perhaps CqsR serves as a dual-function receptor that senses an unknown self-made signal (e.g., autoinducer) and an exogenous metabolic by-product as an environmental cue. A combination of different signals ensures *V. cholerae* only expresses the HCD QS response to repress virulence factor production and biofilm formation at the most appropriate time and environment. This kind of dual detection of cognate AIs and other chemical signals by a single QS receptor is not commonly observed. QseC is one of the known QS receptors that detects both an AI signal (AI-3) and other chemical cues (epinephrine and norepinephrine) to control virulence gene expression in *E. coli* and *Salmonella* (69). It is proposed that such unique QS receptor can be used for communication between the pathogens and the hosts (52). Further work is required to elucidate the role of ethanolamine sensing in inter-kingdom QS signaling and virulence control in *V. cholerae*.

## ACKNOWLEDGEMENTS

We thank James Baleja for assistance with performing NMR metabolomics experiments, and Juan Hernandez and David Giacolone for technical help. WLN, SW, KB, LAH were supported in part by NIH grants AI007329, and AI121337. The funders had no role in study design, data collection and interpretation, or the decision to submit the work for publication

## Detail of chemical synthesis of HEGly and HEHEAA

Unless otherwise noted, all reactions were performed in flame-dried glassware under an atmosphere of nitrogen using dried reagents and solvents. All chemicals were purchased from commercial vendors and used without further purification. Anhydrous solvents were purchased from commercial vendors. Flash chromatography was performed using standard grade silica gel 60 230-400 mesh from SORBENT Technologies or was performed using a Biotage Flash Purification system equipped with Biotage silica gel or C18 columns. Analytical thin-layer chromatography was carried out using Silica G TLC plates, 200 μm with UV_254_ fluorescent indicator (SORBENT Technologies), and visualization was performed by staining and/or by absorbance of UV light. NMR spectra were recorded using a Varian Mercury Plus spectrometer (400 MHz for ^1^H-NMR; 100 MHz for ^13^C-NMR). Chemical shifts are reported in parts per million (ppm) and were calibrated according to residual protonated solvent. Mass spectroscopy data was collected using an Agilent 1100-Series LC/MSD Trap LC-MS or a Micromass Quattromicro with a Waters 2795 Separations Module LC-MS with acetonitrile containing 0.1% formic acid as the mobile phase in positive ionization mode. All final compounds were evaluated to be of greater than 90% purity by analysis of ^1^H-NMR and ^13^C-NMR unless otherwise indicated.

### 2-((2-Hydroxyethyl)amino)acetic Acid, HEGly

This compound was synthesized following the reported procedure (65) with modifications. Briefly, to a solution of ethanolamine (3.15 mL, 52.2 mmol) in anhydrous THF (53mL) in a oven-dried flask was added *i-*Pr_2_NEt (9.4 mL, 54.0 mmol). The resulting solution was cooled to 0°C and was treated with ethyl bromoacetate (5mL, 45.2mmol) dropwise. The mixture was stirred overnight with warming to room temperature and was concentrated *in vacuo* to remove solvent and was purified by silica gel flash chromatography to provide the ester, ethyl (2-hydroxyethyl)glycinate (3.5 g, 23.8 mmol, 53% yield). The ester, ethyl (2-hydroxyethyl)glycinate (3.5g, 23.8 mmol) was dissolved in a solution of acetone (119mL) and water (119mL) and was treated with LiOH (4.99g, 119.01mmol) and stirred at room temperature overnight. The reaction was quenched by the addition of 119mL of 1M HCl and was concentrated to dryness *in vacuo*. The residue was purified by slow recrystallization from H_2_O/EtOH at 5°C to provide 2-((2-Hydroxyethyl)amino)acetic Acid, HEGly (1.10g, 9.2mmol, 39% yield). Spectral data was consistent with previously reported data (65).

### *N*-(2-hydroxyethyl)-2-((2-hydroxyethyl)amino)acetamide, HEHEAA

2-((2-Hydroxyethyl)amino)acetic Acid (HEGly, 87 mg, 0.73 mmol) was dissolved in H_2_O (200 uL) with warming. The solution was diluted with ACN (1.5 mL) and was treated sequentially with *i*-Pr_2_Net (250 uL, 1.46 mmol), 2-aminoethanol (88 uL, 1.46 mmol), 1-Hydroxybenzotriazole (HOBt, 197 mg, 1.46 mmol) and N-(3-Dimethylaminopropyl)-N’-ethylcarbodiimide hydrochloride (EDC, 279 mg, 1.46 mmol). The resulting mixture was heated to 55 °C and was stirred overnight. The reaction mixture was concentrated to dryness and purified using C18 flash chromatography to provide *N*-(2-hydroxyethyl)-2-((2-hydroxyethyl)amino)acetamide, HEHEAA. The product HEHEAA co-eluted with 1-(3-(dimethylamino)propyl)-3-ethylurea and was characterized and used in biological studies as an inseparable ca. 1:1 mixture based on ^1^H-NMR and ^13^C-NMR.

---

**Supplementary Table 1.** List of strains used in this study

| Strain Number | Organism | Genotype | Ab Resistance<sup>a</sup> | Source |
| --- | --- | --- | --- | --- |
| WN3176 | *V. cholerae* | $\Delta cqsS \Delta luxQ \Delta vpsS$ | Sm | (11) |
| WN3208 | *V. cholerae* | $\Delta cqsS \Delta luxQ \Delta vpsS$ / pBB1 | Sm Tet | (11) |
| WN3357 | *V. cholerae* | $\Delta cqsS \Delta luxQ \Delta cqsR$ /pBB1 | Sm Tet | (11) |
| WN3649 | *V. cholerae* | $\Delta cqsS \Delta cqsR \Delta vpsS$ /pBB1 | Sm Tet | (11) |
| WN3651 | *V. cholerae* | $\Delta luxQ \Delta cqsR \Delta vpsS$ /pBB1 | Sm Tet | (11) |
| WN3354 | *V. cholerae* | $\Delta cqsS \Delta luxQ \Delta vpsS \Delta cqsR$ | Sm | (11) |
| WN4899 | *V. cholerae* | $\Delta cqsS \Delta luxQ \Delta vpsS \Delta cqsR$ / pEVS143-*cqsR* / pBB1 | Sm Kan Tet | This study |
| WN5267 | *V. cholerae* | $\Delta cqsS \Delta luxQ \Delta vpsS \Delta cqsR$ / *pEVS143-cqsR*<sup>R49S</sup> / pBB1 | Sm Kan Tet | This study |
| WN5275 | *V. cholerae* | $\Delta cqsS \Delta luxQ \Delta vpsS \Delta cqsR$ / pEVS143-*cqsR*<sup>D198V</sup> / pBB1 | Sm Kan Tet | This study |
| WN5277 | *V. cholerae* | $\Delta cqsS \Delta luxQ \Delta vpsS \Delta cqsR$ / pEVS143-*cqsR*<sup>L217S</sup> / pBB1 | Sm Kan Tet | This study |
| WN5279 | *V. cholerae* | $\Delta cqsS \Delta luxQ \Delta vpsS \Delta cqsR$ / pEVS143-*cqsR*<sup>V219E</sup> / pBB1 | Sm Kan Tet | This study |
| WN5281 | *V. cholerae* | $\Delta cqsS \Delta luxQ \Delta vpsS \Delta cqsR$ / pEVS143-*cqsR*<sup>A259V</sup> / pBB1 | Sm Kan Tet | This study |
| WN5285 | *V. cholerae* | $\Delta cqsS \Delta luxQ \Delta vpsS \Delta cqsR$ / pEVS143-*cqsR*<sup>H262Y</sup> / pBB1 | Sm Kan Tet | This study |
| WN5318 | *V. cholerae* | $\Delta cqsS \Delta luxQ \Delta vpsS \Delta cqsR$ / pEVS143-*cqsR*<sup>F268I</sup> / pBB1 | Sm Kan Tet | This study |
| WN5354 | *V. cholerae* | $\Delta cqsS \Delta luxQ \Delta vpsS \Delta cqsR$ / pEVS143-*cqsR*<sup>D171V</sup> / pBB1 | Sm Kan Tet | This study |
| WN5781 | *V. cholerae* | $\Delta cqsS \Delta luxQ \Delta vpsS \Delta vc1554 \Delta vca0136$ | Sm | This study |
| WN5784 | *V. cholerae* | $\Delta cqsS \Delta luxQ \Delta vpsS \Delta vc1554 \Delta vca0136$/pBB1 | Sm Tet | This study |
| WN5886 | *V. cholerae* | $\Delta cqsS \Delta luxQ \Delta vpsS cqsR$<sup>D198V</sup> | Sm | This study |
| WN5887 | *V. cholerae* | $\Delta cqsS \Delta luxQ \Delta vpsS cqsR$<sup>D171V</sup> | Sm | This study |
| WN5890 | *V. cholerae* | $\Delta cqsS \Delta luxQ \Delta vpsS cqsR$<sup>D198V</sup>/pBB1 | Sm Tet | This study |
| WN5891 | *V. cholerae* | $\Delta cqsS \Delta luxQ \Delta vpsS cqsR$<sup>D171V</sup>/pBB1 | Sm Tet | This study |
| WN5976 | *V. cholerae* | $\Delta cqsS \Delta luxQ \Delta vpsS$ /pBK1003 (*Pqrr4-lux*) | Sm Cm | This study |
| WN5979 | *V. cholerae* | $\Delta cqsS \Delta luxQ \Delta vpsS \Delta vc1554 \Delta vca0136$ / pBK1003 | Sm Cm | This study |
| WN5980 | *V. cholerae* | $\Delta cqsS \Delta luxQ \Delta vpsS cqsR$<sup>D198V</sup> / pBK1003 | Sm Cm | This study |
| WN5981 | *V. cholerae* | $\Delta cqsS \Delta luxQ \Delta vpsS cqsR$<sup>D171V</sup> / pBK1003 | Sm Cm | This study |
| WN048 | *E. coli* | *S17-1* λpir / pBB1 | Tet | (11) |
| WN3705 | *E. coli* | *S17-1* λpir / pEVS143-*cqsR* | Kan | (11) |
| WN5233 | *E. coli* | XL10-Gold / *pEVS143-cqsR*<sup>V219E</sup> | Kan | This study |
| WN5236 | *E. coli* | XL10-Gold / *pEVS143-cqsR*<sup>A259V</sup> | Kan | This study |
| WN5239 | *E. coli* | XL10-Gold / *pEVS143-cqsR*<sup>H262Y</sup> | Kan | This study |
| **WN5242** | *E. coli* | XL10-Gold / *pEVS143-cqsR*<sup>R49S</sup> | Kan | This study |
| **WN5248** | *E. coli* | XL10-Gold / *pEVS143-cqsR*<sup>D198V</sup> | Kan | This study |
| **WN5250** | *E. coli* | XL10-Gold / *pEVS143-cqsR*<sup>L217S</sup> | Kan | This study |
| **WN5318** | *E. coli* | XL10-Gold / *pEVS143-cqsR*<sup>F268I</sup> | Kan | This study |
| **WN5326** | *E. coli* | BL21DE3/ pET28b *cqsR* -LBD | Kan | This study |
| **WN5344** | *E. coli* | XL10-Gold / *pEVS143-cqsR*<sup>D171V</sup> | Kan | This study |
| **WN5745** | *E. coli* | S17 λpir DAP- / pKAS::Kan *Δvc1554* | Kan | This study |
| **WN5746** | *E. coli* | S17 λpir DAP- / pKAS::Kan *Δvca0136* | Kan | This study |
| **WN5880** | *E. coli* | S17 λpir DAP- / pKAS::Kan *cqsR*<sup>D198V</sup> | Kan | This study |
| **WN5881** | *E. coli* | S17 λpir DAP- / pKAS::Kan *cqsR*<sup>D171V</sup> | Kan | This study |

<sup>a</sup> Sm = Streptomycin, Kan = Kanamycin, Cm = Chloramphenicol, Tet = Tetracycline

**Supplementary Table 2.** Binding activity of CqsR to common byproducts of ethanolamine

Binding activity of CqsR to common byproducts of ethanolamine
| Compound | T<sub>m</sub><sup>a</sup> |
| --- | --- |
| None | 47.00 |
| N-(2-hydroxyethyl)-acetamide | 48.00 |
| N,N'-Bis(2-hydroxyethyl)-ethanediamide | 49.00 |
| N-(2-hydroxyethyl)-2-[(2-hydroxyethyl)amino]-acetamide | 47.67 [0.57] |
| N-(2-hydroxyethyl)ethylenediamine | 47.67 [0.57] |
| N-(2-hydroxyethyl) imidazolidone | 47.00 |
| N-(2-hydroxyethyl)-glycine | 47.33 [0.57] |
| N-(2-hydroxyethyl)-imidazole | 47.33 [0.57] |
| N-(2-hydroxyethyl)-formamide | 50.00 |

<sup>a</sup>Data shown as average of 3 replicates in the presence of 1 mM tested compounds. Square brackets indicate standard deviation where applicable.

**Supplementary Figure 1.**
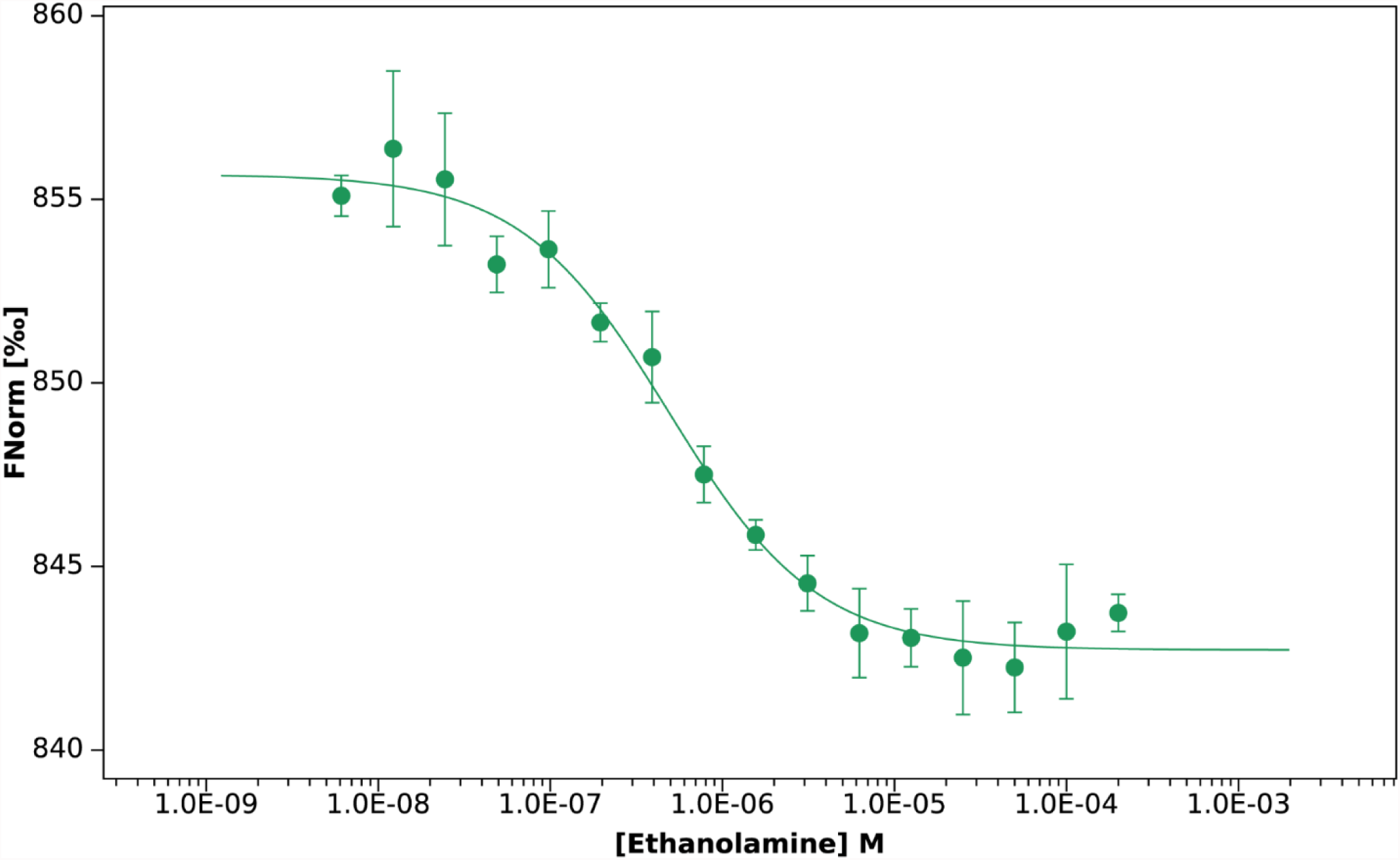
MST quantification (FNorm; normalized fluorescence) for ethanolamine binding to CqsR. Ethanolamine was titrated between 200 μM and 0.0061 μM with 20 nM His6-tagged CqsR. Ethanolamine binds to CqsR with a K_d_ of 0.478 ± 0.076 µM. Binding affinity was calculated from three independent experiments.

**Supplementary Figure 2.**
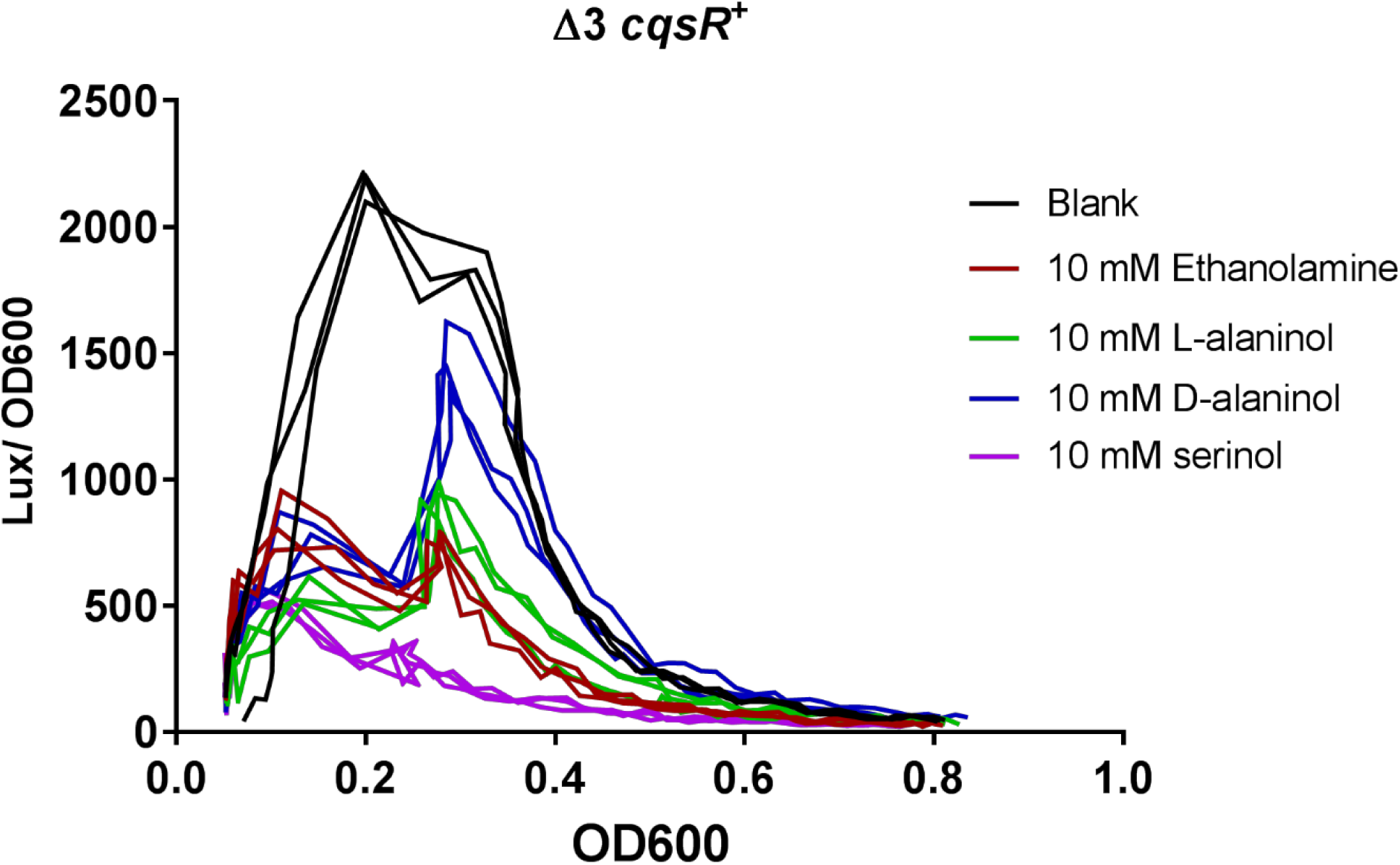
Effect of ethanolamine and its analogs on *qrr*4 transcription. Normalized bioluminescence production (lux/OD_600_) using a P*qrr*4-*lux* reporter was measured in a Δ3 *cqsR*^+^ strain in the presence of 10 mM ethanolamine, L-alaninol, D-alaninol, or serinol. Blank indicates LB medium without added compound.

**Supplementary Figure 3.**
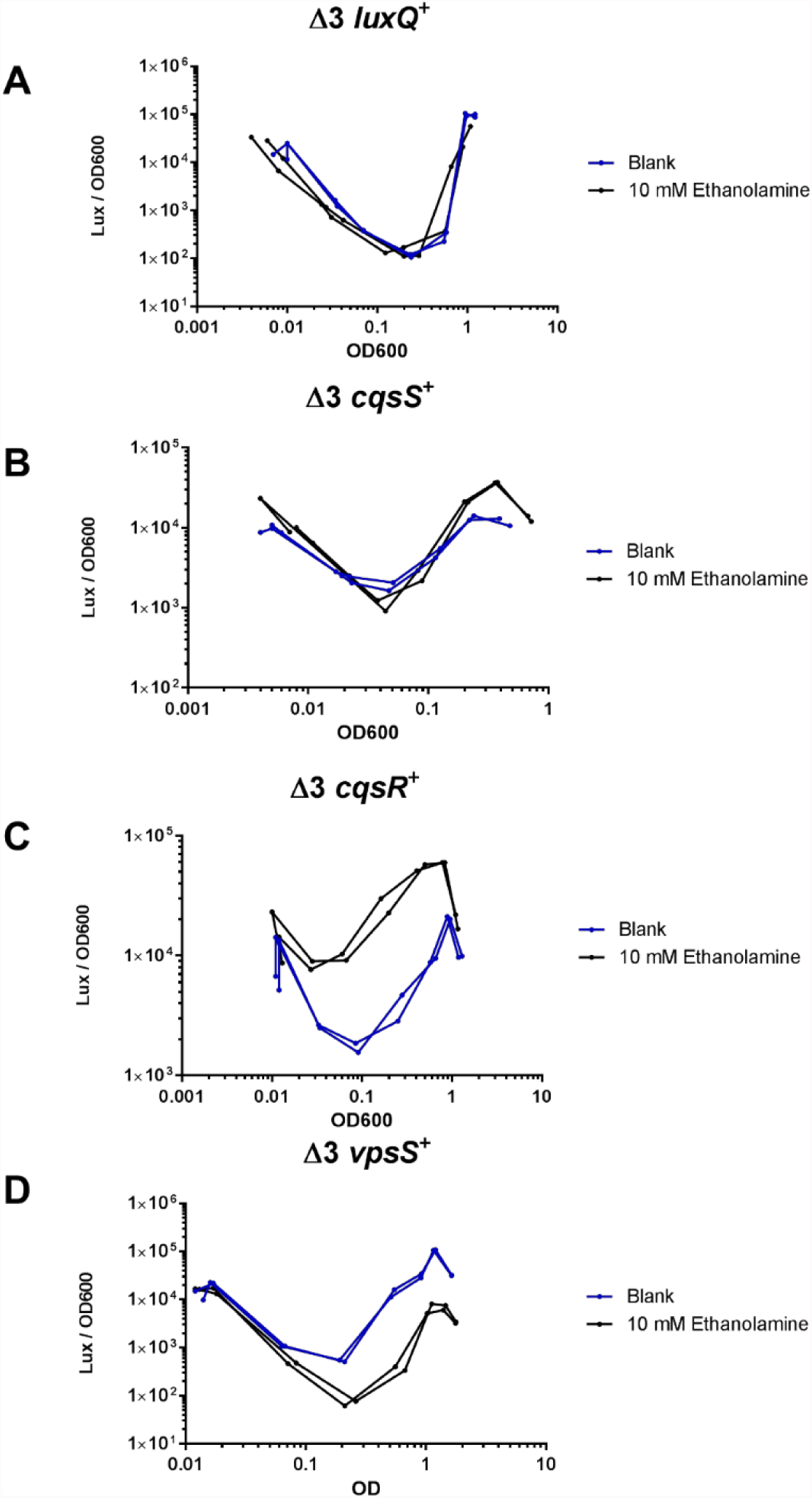
Ethanolamine induces high cell density QS response in a CqsR-specific manner. HapR-dependent bioluminescence profiles (lux/OD_600_) were measured in A) Δ3 *luxQ*^+^, B) Δ3 *cqsS*^+^, C) Δ3 *cqsR*^+^, and D) Δ3 *vpsS*^+^, in LB medium and LB medium containing 10 mM ethanolamine. Each figure shows a representative profile of each condition with two biological replicates. Each experiment was performed independently at least two times.

**Supplementary Figure 4.**
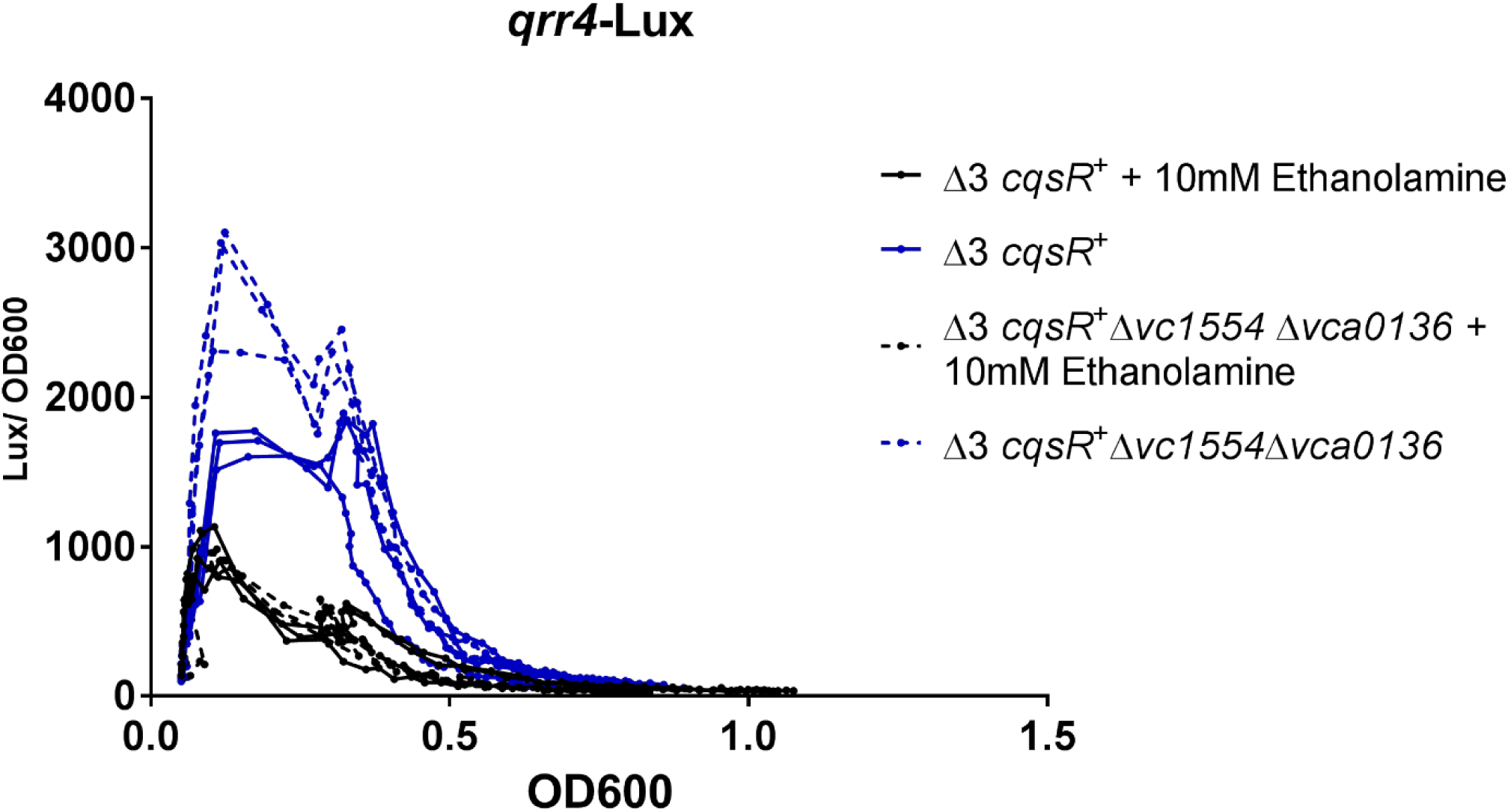
Effect of ethanolamine on *qrr*4 transcription in the double phosphodiesterase mutants. Normalized bioluminescence production (lux/OD_600_) using a P*qrr*4-*lux* reporter was measured in a Δ3 *cqsR*^+^ and the isogenic Δ3 *cqsR* ^+^ Δ*vc1554* Δ*vca0136* strains with and without 10 mM ethanolamine. Each figure shows a representative profile of each condition with three biological replicates. Each experiment was performed independently at least two times.

## REFERENCES

1. Hawver LA, Jung SA, Ng WL. 2016. Specificity and complexity in bacterial quorum-sensing systems. FEMS Microbiol Rev 40:738–52.

2. Suckow G, Seitz P, Blokesch M. 2011. Quorum sensing contributes to natural transformation of *Vibrio cholerae* in a species-specific manner. J Bacteriol 193:4914–24.

3. Lo Scrudato M, Blokesch M. 2012. The regulatory network of natural competence and transformation of *Vibrio cholerae*. PLoS Genet 8:e1002778.

4. Hammer BK, Bassler BL. 2003. Quorum sensing controls biofilm formation in *Vibrio cholerae*. Mol Microbiol 50:101–4.

5. Miller MB, Skorupski K, Lenz DH, Taylor RK, Bassler BL. 2002. Parallel quorum sensing systems converge to regulate virulence in *Vibrio cholerae*. Cell 110:303–14.

6. Zhu J, Miller MB, Vance RE, Dziejman M, Bassler BL, Mekalanos JJ. 2002. Quorum-sensing regulators control virulence gene expression in *Vibrio cholerae*. Proc Natl Acad Sci U S A 99:3129–34.

7. Shao Y, Bassler BL. 2014. Quorum regulatory small RNAs repress type VI secretion in *Vibrio cholerae*. Mol Microbiol 92:921–30.

8. Hawver LA, Giulietti JM, Baleja JD, Ng WL. 2016. Quorum Sensing Coordinates Cooperative Expression of Pyruvate Metabolism Genes To Maintain a Sustainable Environment for Population Stability. MBio 7.

9. Beyhan S, Bilecen K, Salama SR, Casper-Lindley C, Yildiz FH. 2007. Regulation of rugosity and biofilm formation in Vibrio cholerae: comparison of VpsT and VpsR regulons and epistasis analysis of vpsT, vpsR, and hapR. J Bacteriol 189:388–402.

10. Shikuma NJ, Fong JC, Odell LS, Perchuk BS, Laub MT, Yildiz FH. 2009. Overexpression of VpsS, a hybrid sensor kinase, enhances biofilm formation in *Vibrio cholerae*. J Bacteriol 191:5147–58.

11. Jung SA, Chapman CA, Ng WL. 2015. Quadruple quorum-sensing inputs control Vibrio cholerae virulence and maintain system robustness. PLoS Pathog 11:e1004837.

12. Rutherford ST, van Kessel JC, Shao Y, Bassler BL. 2011. AphA and LuxR/HapR reciprocally control quorum sensing in vibrios. Genes Dev 25:397–408.

13. Shao Y, Bassler BL. 2012. Quorum-sensing non-coding small RNAs use unique pairing regions to differentially control mRNA targets. Mol Microbiol 83:599–611.

14. Bardill JP, Zhao X, Hammer BK. 2011. The Vibrio cholerae quorum sensing response is mediated by Hfq-dependent sRNA/mRNA base pairing interactions. Mol Microbiol 80:1381–94.

15. Lenz DH, Mok KC, Lilley BN, Kulkarni RV, Wingreen NS, Bassler BL. 2004. The small RNA chaperone Hfq and multiple small RNAs control quorum sensing in *Vibrio harveyi* and *Vibrio cholerae*. Cell 118:69–82.

16. Higgins DA, Pomianek ME, Kraml CM, Taylor RK, Semmelhack MF, Bassler BL. 2007. The major *Vibrio cholerae* autoinducer and its role in virulence factor production. Nature 450:883–6.

17. Ng WL, Wei Y, Perez LJ, Cong J, Long T, Koch M, Semmelhack MF, Wingreen NS, Bassler BL. 2010. Probing bacterial transmembrane histidine kinase receptor-ligand interactions with natural and synthetic molecules. Proc Natl Acad Sci U S A 107:5575–80.

18. Ng WL, Perez LJ, Wei Y, Kraml C, Semmelhack MF, Bassler BL. 2011. Signal production and detection specificity in Vibrio CqsA/CqsS quorum-sensing systems. Mol Microbiol 79:1407–17.

19. Wei Y, Perez LJ, Ng WL, Semmelhack MF, Bassler BL. 2011. Mechanism of *Vibrio cholerae* Autoinducer-1 Biosynthesis. ACS Chem Biol 6:356–365.

20. Wei Y, Ng WL, Cong J, Bassler BL. 2012. Ligand and antagonist driven regulation of the *Vibrio cholerae* quorum-sensing receptor CqsS. Mol Microbiol 83:1095–108.

21. Surette MG, Miller MB, Bassler BL. 1999. Quorum sensing in *Escherichia coli*, *Salmonella typhimurium*, and *Vibrio harveyi*: a new family of genes responsible for autoinducer production. Proc Natl Acad Sci U S A 96:1639–44.

22. Schauder S, Shokat K, Surette MG, Bassler BL. 2001. The LuxS family of bacterial autoinducers: biosynthesis of a novel quorum-sensing signal molecule. Mol Microbiol 41:463–76.

23. Chen X, Schauder S, Potier N, Van Dorsselaer A, Pelczer I, Bassler BL, Hughson FM. 2002. Structural identification of a bacterial quorum-sensing signal containing boron. Nature 415:545–9.

24. Neiditch MB, Federle MJ, Miller ST, Bassler BL, Hughson FM. 2005. Regulation of LuxPQ receptor activity by the quorum-sensing signal autoinducer-2. Mol Cell 18:507–18.

25. Neiditch MB, Federle MJ, Pompeani AJ, Kelly RC, Swem DL, Jeffrey PD, Bassler BL, Hughson FM. 2006. Ligand-induced asymmetry in histidine sensor kinase complex regulates quorum sensing. Cell 126:1095–108.

26. Papenfort K, Silpe JE, Schramma KR, Cong JP, Seyedsayamdost MR, Bassler BL. 2017. A Vibrio cholerae autoinducer-receptor pair that controls biofilm formation. Nat Chem Biol 13:551–557.

27. Hossain S, Heckler I, Boon EM. 2018. Discovery of a Nitric Oxide Responsive Quorum Sensing Circuit in Vibrio cholerae. ACS Chem Biol 13:1964–1969.

28. Upadhyay AA, Fleetwood AD, Adebali O, Finn RD, Zhulin IB. 2016. Cache Domains That are Homologous to, but Different from PAS Domains Comprise the Largest Superfamily of Extracellular Sensors in Prokaryotes. PLoS Comput Biol 12:e1004862.

29. Marchler-Bauer A, Derbyshire MK, Gonzales NR, Lu S, Chitsaz F, Geer LY, Geer RC, He J, Gwadz M, Hurwitz DI, Lanczycki CJ, Lu F, Marchler GH, Song JS, Thanki N, Wang Z, Yamashita RA, Zhang D, Zheng C, Bryant SH. 2015. CDD: NCBI’s conserved domain database. Nucleic Acids Res 43:D222–6.

30. Geer LY, Domrachev M, Lipman DJ, Bryant SH. 2002. CDART: protein homology by domain architecture. Genome Res 12:1619–23.

31. Mitchell AL, Attwood TK, Babbitt PC, Blum M, Bork P, Bridge A, Brown SD, Chang HY, El-Gebali S, Fraser MI, Gough J, Haft DR, Huang H, Letunic I, Lopez R, Luciani A, Madeira F, Marchler-Bauer A, Mi H, Natale DA, Necci M, Nuka G, Orengo C, Pandurangan AP, Paysan-Lafosse T, Pesseat S, Potter SC, Qureshi MA, Rawlings ND, Redaschi N, Richardson LJ, Rivoire C, Salazar GA, Sangrador-Vegas A, Sigrist CJA, Sillitoe I, Sutton GG, Thanki N, Thomas PD, Tosatto SCE, Yong SY, Finn RD. 2019. InterPro in 2019: improving coverage, classification and access to protein sequence annotations. Nucleic Acids Res 47:D351–D360.

32. Krogh A, Larsson B, von Heijne G, Sonnhammer EL. 2001. Predicting transmembrane protein topology with a hidden Markov model: application to complete genomes. J Mol Biol 305:567–80.

33. Zimmermann L, Stephens A, Nam SZ, Rau D, Kubler J, Lozajic M, Gabler F, Soding J, Lupas AN, Alva V. 2018. A Completely Reimplemented MPI Bioinformatics Toolkit with a New HHpred Server at its Core. Journal of Molecular Biology 430:2237–2243.

34. Kelley LA, Mezulis S, Yates CM, Wass MN, Sternberg MJE. 2015. The Phyre2 web portal for protein modeling, prediction and analysis. Nature Protocols 10:845–858.

35. Nishiyama S, Takahashi Y, Yamamoto K, Suzuki D, Itoh Y, Sumita K, Uchida Y, Homma M, Imada K, Kawagishi I. 2016. Identification of a Vibrio cholerae chemoreceptor that senses taurine and amino acids as attractants. Sci Rep 6:20866.

36. Thelin KH, Taylor RK. 1996. Toxin-coregulated pilus, but not mannose-sensitive hemagglutinin, is required for colonization by *Vibrio cholerae* O1 El Tor biotype and O139 strains. Infect Immun 64:2853–6.

37. Skorupski K, Taylor RK. 1996. Positive selection vectors for allelic exchange. Gene 169:47–52.

38. Thomas J, Watve SS, Ratcliff WC, Hammer BK. 2017. Horizontal Gene Transfer of Functional Type VI Killing Genes by Natural Transformation. MBio 8.

39. Bose JL, Rosenberg CS, Stabb EV. 2008. Effects of luxCDABEG induction in Vibrio fischeri: enhancement of symbiotic colonization and conditional attenuation of growth in culture. Arch Microbiol 190:169–83.

40. McKellar JL, Minnell JJ, Gerth ML. 2015. A high-throughput screen for ligand binding reveals the specificities of three amino acid chemoreceptors from Pseudomonas syringae pv. actinidiae. Mol Microbiol 96:694–707.

41. Anantharaman V, Aravind L. 2000. Cache - a signaling domain common to animal Ca(2+)-channel subunits and a class of prokaryotic chemotaxis receptors. Trends Biochem Sci 25:535–7.

42. Coutinho BG, Mevers E, Schaefer AL, Pelletier DA, Harwood CS, Clardy J, Greenberg EP. 2018. A plant-responsive bacterial-signaling system senses an ethanolamine derivative. Proc Natl Acad Sci U S A 115:9785–9790.

43. Vevelstad SJ, Johansen MT, Knuutila H, Svendsen HF. 2016. Extensive dataset for oxidative degradation of ethanolamine at 55–75°C and oxygen concentrations from 6 to 98%. International Journal of Greenhouse Gas Control 50:158–178.

44. Ohshima N, Yamashita S, Takahashi N, Kuroishi C, Shiro Y, Takio K. 2008. Escherichia coli cytosolic glycerophosphodiester phosphodiesterase (UgpQ) requires Mg2+, Co2+, or Mn2+ for its enzyme activity. J Bacteriol 190:1219–23.

45. Tommassen J, Eiglmeier K, Cole ST, Overduin P, Larson TJ, Boos W. 1991. Characterization of two genes, glpQ and ugpQ, encoding glycerophosphoryl diester phosphodiesterases of Escherichia coli. Mol Gen Genet 226:321–7.

46. Brewster JL, McKellar JL, Finn TJ, Newman J, Peat TS, Gerth ML. 2016. Structural basis for ligand recognition by a Cache chemosensory domain that mediates carboxylate sensing in Pseudomonas syringae. Sci Rep 6:35198.

47. Gavira JA, Ortega A, Martin-Mora D, Conejero-Muriel MT, Corral-Lugo A, Morel B, Matilla MA, Krell T. 2018. Structural Basis for Polyamine Binding at the dCACHE Domain of the McpU Chemoreceptor from Pseudomonas putida. J Mol Biol 430:1950–1963.

48. Day CJ, King RM, Shewell LK, Tram G, Najnin T, Hartley-Tassell LE, Wilson JC, Fleetwood AD, Zhulin IB, Korolik V. 2016. A direct-sensing galactose chemoreceptor recently evolved in invasive strains of Campylobacter jejuni. Nat Commun 7:13206.

49. Shrestha M, Compton KK, Mancl JM, Webb BA, Brown AM, Scharf BE, Schubot FD. 2018. Structure of the sensory domain of McpX from Sinorhizobium meliloti, the first known bacterial chemotactic sensor for quaternary ammonium compounds. Biochem J 475:3949–3962.

50. Webb BA, Karl Compton K, Castaneda Saldana R, Arapov TD, Keith Ray W, Helm RF, Scharf BE. 2017. Sinorhizobium meliloti chemotaxis to quaternary ammonium compounds is mediated by the chemoreceptor McpX. Mol Microbiol 103:333–346.

51. Kendall MM, Gruber CC, Parker CT, Sperandio V. 2012. Ethanolamine controls expression of genes encoding components involved in interkingdom signaling and virulence in enterohemorrhagic Escherichia coli O157:H7. MBio 3.

52. Del Papa MF, Perego M. 2008. Ethanolamine activates a sensor histidine kinase regulating its utilization in Enterococcus faecalis. J Bacteriol 190:7147–56.

53. Martin-Mora D, Ortega A, Perez-Maldonado FJ, Krell T, Matilla MA. 2018. The activity of the C4-dicarboxylic acid chemoreceptor of Pseudomonas aeruginosa is controlled by chemoattractants and antagonists. Sci Rep 8:2102.

54. Khatri N, Khatri I, Subramanian S, Raychaudhuri S. 2012. Ethanolamine utilization in Vibrio alginolyticus. Biol Direct 7:45; discussion 45.

55. Penrod JT, Mace CC, Roth JR. 2004. A pH-sensitive function and phenotype: evidence that EutH facilitates diffusion of uncharged ethanolamine in Salmonella enterica. J Bacteriol 186:6885–90.

56. Garsin DA. 2010. Ethanolamine utilization in bacterial pathogens: roles and regulation. Nat Rev Microbiol 8:290–5.

57. Anderson CJ, Clark DE, Adli M, Kendall MM. 2015. Ethanolamine Signaling Promotes Salmonella Niche Recognition and Adaptation during Infection. PLoS Pathog 11:e1005278.

58. Kaval KG, Garsin DA. 2018. Ethanolamine Utilization in Bacteria. MBio 9.

59. Nawrocki KL, Wetzel D, Jones JB, Woods EC, McBride SM. 2018. Ethanolamine is a valuable nutrient source that impacts Clostridium difficile pathogenesis. 20:1419–1435.

60. Bertin Y, Girardeau JP, Chaucheyras-Durand F, Lyan B, Pujos-Guillot E, Harel J, Martin C. 2011. Enterohaemorrhagic Escherichia coli gains a competitive advantage by using ethanolamine as a nitrogen source in the bovine intestinal content. Environ Microbiol 13:365–77.

61. Anderson CJ, Kendall MM. 2016. Location, location, location. Salmonella senses ethanolamine to gauge distinct host environments and coordinate gene expression. Microb Cell 3:89–91.

62. Thiennimitr P, Winter SE, Winter MG, Xavier MN, Tolstikov V, Huseby DL, Sterzenbach T, Tsolis RM, Roth JR, Baumler AJ. 2011. Intestinal inflammation allows Salmonella to use ethanolamine to compete with the microbiota. Proc Natl Acad Sci U S A 108:17480–5.

63. Federle MJ, Bassler BL. 2003. Interspecies communication in bacteria. J Clin Invest 112:1291–9.

64. Pereira CS, Thompson JA, Xavier KB. 2013. AI-2-mediated signalling in bacteria. FEMS Microbiol Rev 37:156–81.

65. Ashley JD, Stefanick JF, Schroeder VA, Suckow MA, Kiziltepe T, Bilgicer B. 2014. Liposomal bortezomib nanoparticles via boronic ester prodrug formulation for improved therapeutic efficacy in vivo. J Med Chem 57:5282–92.

